# Redefining the role of the *Plasmodium* heme detoxification protein: From hemozoin formation to mitochondrial protein synthesis

**DOI:** 10.1101/2025.11.27.690934

**Authors:** Louis Sarrazin, Melissa R. Rosenthal, Joachim Kloehn, Tanja Ziesmann, Yasmin Schmitz, Anna-Lena Sandtmann, Robert Domènech-Eres, Katharina Scholz-Höhn, Coralie Boulet, Ute Distler, Daniel E. Goldberg, Joachim M. Matz

## Abstract

Throughout their intraerythrocytic development, malaria parasites digest up to 80% of the host cell’s hemoglobin within a specialized degradative compartment known as the digestive vacuole. This process releases heme, which is detoxified by sequestration into bioinert hemozoin crystals. Although heme biomineralization is essential for blood-stage survival and a validated drug target, its underlying mechanisms remain unclear. Initially identified as a potent inducer of β-hematin crystallization *in vitro*, the parasite’s <u>H</u>eme <u>D</u>etoxification <u>P</u>rotein (HDP) has been proposed to execute a similar role in the formation of hemozoin crystals *in cellulo*. Here, we investigate the function of HDP in live *Plasmodium falciparum* parasites, integrating experimental genetic approaches with quantitative microscopy, cellular bioenergetics and whole-proteome profiling. Endogenous tagging revealed that HDP localizes to the mitochondrion rather than the digestive vacuole. Conditional inactivation of HDP resulted in a gradual loss of mitochondrial membrane potential, preceding developmental arrest. Bypassing the essential role of the respiratory chain in pyrimidine biosynthesis – either through exogenous electron acceptors or expression of a ubiquinone-independent dihydroorotate dehydrogenase – rescued HDP-deficient parasites, indicating a role in maintaining respiratory chain activity. Consistent with this, electron flow through complex IV was abolished in rescued HDP-null parasites, rendering them hypersensitive to proguanil, an antimalarial that synergizes with respiratory chain inhibitors. We found that loss of HDP leads to a marked reduction of complexes III and IV, whose integrity depends on mitochondrial protein biosynthesis. Integration of quantitative proteomic data with structure-guided homology modelling supports a role for HDP as part of the large mitoribosomal subunit at the inter-subunit contact site. By contrast, HDP loss did not affect the quantity of hemozoin or other heme species, crystal morphology, or sensitivity to the hemozoin-targeting drug chloroquine. Together, these findings challenge previous models linking HDP to hemozoin formation and instead reveal an essential role for HDP in mitochondrial protein biosynthesis.

## INTRODUCTION

Heme is a ubiquitous and evolutionarily ancient prosthetic group that plays a central role in a wide range of biological processes. It consists of a porphyrin ring coordinating a central iron atom (1), which can alternate between ferrous (Fe^2+^) and ferric (Fe^3+^) states, allowing it to accept and donate electrons in redox reactions and to bind small gaseous molecules reversibly. Heme-containing proteins include cytochromes, which facilitate electron transfer during photosynthesis and respiration, and globins, which bind gases for transport, storage, and sensing (2, 3). In addition, various enzymes use heme iron as a catalytic center to carry out diverse redox or radical reactions (4).

The limited bioavailability of iron and the high metabolic cost of heme biosynthesis can constrain growth and development. In pathogens, this constraint is amplified during infection, when hosts sequester heme and iron through nutritional immunity, which in turn drives the evolution of specialized acquisition strategies such as high affinity transport systems, siderophore-based uptake pathways, and other mechanisms that enable pathogens to compensate for host imposed nutrient limitation (5). In this regard, malaria parasites differ markedly from most pathogens. These vector-borne protozoans replicate within red blood cells (RBCs), an environment exceptionally rich in heme (6). Although the parasite genome encodes a heme biosynthetic pathway, it is dispensable during blood-stage development and only becomes essential later in the mosquito vector and the human liver, where heme is less abundant (7–9).

Asexual blood-stage parasites nevertheless rely on heme-mediated reactions, particularly in the mitochondrial electron transport chain (ETC) (10). Their ETC is highly unusual among eukaryotes, as it is not required for ATP synthesis. Components of the ATP synthase (complex V) and enzymes of the tricarboxylic acid (TCA) cycle are dispensable at this stage, with most ATP generated via glycolysis (11–16). Nonetheless, the mitochondrion maintains an active ETC, in which electrons are sequentially passed along heme-containing cytochromes before ultimately reducing oxygen (17). This electron flow powers proton pumping by respiratory complexes III and IV, thereby generating a chemiosmotic gradient that helps sustain the mitochondrial membrane potential (ΔΨ_mt_) (18).

The integrity of complexes III and IV depends on both cytosolic and mitochondrial protein synthesis. While most subunits are nuclear encoded and imported into the mitochondrion post-translationally, three proteins are encoded in the mitochondrial genome: cytochrome b of complex III, and cytochrome c oxidase subunits 1 (Cox1) and 3 (Cox3) of complex IV (19, 20). In apicomplexan parasites, these are the only proteins synthesized by the mitochondrial ribosome (mitoribosome). The mitoribosome is a large complex composed of highly fragmented rRNAs and both conserved as well as parasite-specific proteins, all organized into a large (LSU) and a small subunit (SSU), functionally streamlined to produce only these three proteins (21, 22).

Complex III receives electrons from the lipid-soluble carrier ubiquinol, a transfer that is essential for parasite survival, as illustrated by the antimalarial drug atovaquone, which blocks this reaction (23, 24). In most systems, ubiquinol is generated by dehydrogenases of the TCA cycle and by complex I which uses NADH to reduce ubiquinone. However, complex I is absent in malaria parasites. They instead possess an alternative single-subunit NADH:ubiquinone oxidoreductase, which is not required for blood-stage growth, similar to the dehydrogenases of the TCA cycle (11, 25, 26). This suggests that their contribution to feeding electrons into complex III is minor. The main enzyme donating electrons to complex III is believed to be dihydroorotate dehydrogenase (DHODH), which produces the pyrimidine precursor orotate and transfers the electrons to ubiquinone (10). Interestingly, the only essential role of complexes III and IV during asexual blood-stage development is to enable pyrimidine biosynthesis by serving as an electron sink that regenerates ubiquinone for DHODH activity (10). Bypassing this requirement, either by supplying alternative electron acceptors or by expressing a ubiquinone-independent DHODH, fully restores parasite viability when the terminal respiratory complexes are pharmacologically or genetically disrupted (10, 27).

The heme that enables electron transfer within the ETC is salvaged from the host RBC, whose soluble protein content is >95% hemoglobin (7–9). Hemoglobin is endocytosed and transported to a lysosome-like organelle known as the digestive vacuole (DV), where it is proteolytically degraded, providing the parasite with amino acids and releasing heme for potential use (28). However, far more heme is mobilized than needed. Owing to its reactive nature, free heme iron is highly cytotoxic, damaging lipids, proteins and DNA (29, 30). Consequently, the parasite must balance its metabolic requirement for heme with an effective detoxification strategy. This is achieved by sequestering the vast majority of heme within the DV as bioinert hemozoin crystals (30, 31). Upon release from the globin chains, heme is first oxidized to hematin, which forms dimers through reciprocal iron-carboxylate bonds. These dimers then polymerize via hydrogen bonding to form the hemozoin crystals (32). Heme biomineralization is essential for parasite survival and is targeted by potent antimalarials, such as chloroquine, which inhibit adsorption of hematin onto the growing crystal surface (33–36).

The mechanism by which heme forms hemozoin remains debated. Minimalistic *in vitro* assays have been used to test how various additives affect the formation of β-hematin, a synthetic but structurally identical form of hemozoin. These studies have produced conflicting results, implicating lipids (37, 38), membranes (39), phase boundaries (40), and proteins (41, 42) as potential drivers of heme biomineralization. A few parasite proteins have been shown to modulate heme crystallization *in vitro*, and one even *in cellulo,* yet none appear essential for parasite survival (41, 43). The notable exception is the heme detoxification protein (HDP, PF3D7_1446800), which efficiently induces β-hematin formation *in vitro* and for which genetic evidence supports a critical role in parasite viability (14, 15, 42). Biochemical and imaging-based studies also indicate that HDP may reside in the DV, supporting a potential function in heme biomineralization (42, 44, 45). However, parasite-centered research approaches have yet to demonstrate this role *in cellulo*.

Here, we characterize the function of HDP in the human malaria pathogen *Plasmodium falciparum* using reverse genetic approaches. We show that HDP is not required for hemozoin formation, but instead participates in mitochondrial protein biosynthesis, likely as a component of the mitoribosome, which is essential for the integrity and function of the ETC. These findings represent a paradigm shift in the study of heme biomineralization, revealing an alternative role for a protein long considered the primary driver of hemozoin formation.

## RESULTS

### HDP localizes to the parasite mitochondrion

Previous studies have reported conflicting localizations for HDP, placing the protein in the RBC cytosol, the DV, and elongated organelles within the parasite (42, 45). To clarify its precise subcellular distribution, we endogenously fused mNeonGreen (mNG) to the C-terminus of HDP, allowing direct visualization in live parasites. Simultaneously, we introduced two *loxP* sites into the HDP locus, one within the first intron and another downstream of the stop codon, to enable DiCre-mediated conditional reverse genetic experiments (46) (Figure 1A, Supplementary Table S1). Following transfection of the modifying construct into DiCre-expressing 3D7 parasites (strain: B11) (47), two *hdp-mNG:loxP* clones were isolated, and correct genomic integration was confirmed by diagnostic PCR (Figure 1B).

**Figure 1:**
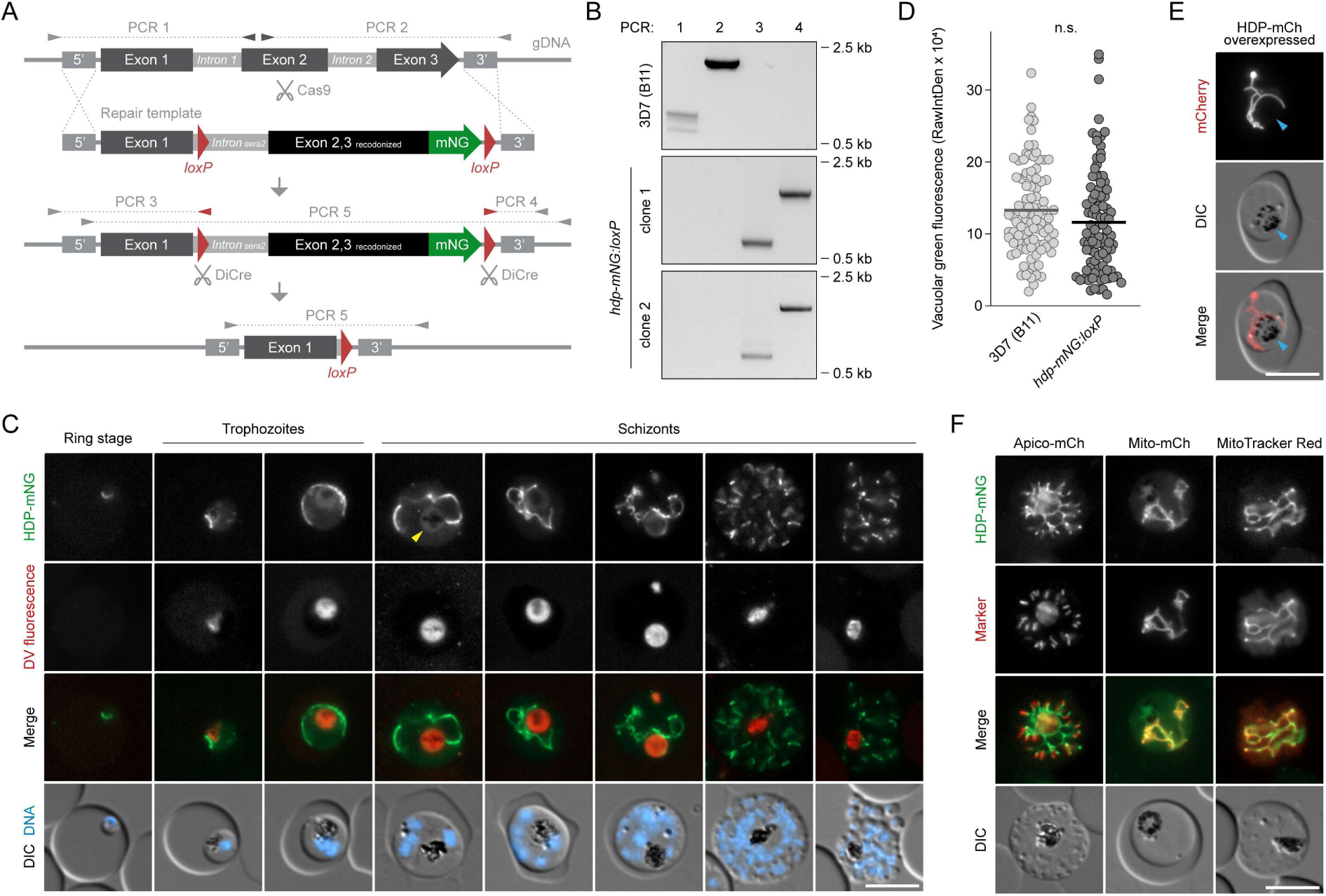
HDP localizes to the parasite mitochondrion. **(A)** Genetic strategy for tagging and conditional deletion of HDP using Cas9 and the DiCre system. Shown are the endogenous locus, the synthetic repair template, and the recombined locus before and after excision. Primer combinations for diagnostic PCRs are indicated. **(B)** Diagnostic PCRs as in (A), using genomic DNA isolated from the parental line and two independent *hdp-mNG:loxP* clones. **(C)** Expression and localization of HDP-mNG observed by live fluorescence microscopy. Representative images are shown throughout the intraerythrocytic cycle. The DV was visualized via porphyrin-derived autofluorescence. Yellow arrowhead: faint signal in the DV. **(D)** Green DV fluorescence in *hdp-mNG:loxP* parasites does not exceed the background observed in parental parasites 36 hours post-invasion. Shown are mean and individual values. n.s., non-significant, paired t-test; N = 99 parasites from 3 independent experiments. **(E)** Overexpressed HDP-mCh localizes exclusively to a branching organelle. Blue arrowheads: position of the DV. **(F)** HDP-mNG colocalizes with mitochondrial markers. Shown are *hdp-mNG:loxP* parasites expressing fluorescent organelle markers (Apico-mCh, Mito-mCh) or stained with MitoTracker Red CMXRos. Scale bars, 5 µm.

Live fluorescence microscopy revealed mNG signal in a tubular parasite organelle, which expanded dramatically during parasite maturation, and eventually branched and divided during merozoite segmentation (Figure 1C). In addition, a faint green signal colocalized with the porphyrin-derived red autofluorescence of the DV, but comparison with parental B11 parasites revealed that it did not exceed the DV’s baseline fluorescence (Figure 1C,D). To further examine a potential DV localization, we overexpressed HDP fused to mCherry (mCh), but despite the much stronger fluorescence, localization remained restricted to the branching organelle, with no apparent signal in the DV (Figure 1E).

The branching morphology of the HDP-containing compartment was reminiscent of the parasite’s two endosymbiotic organelles: the apicoplast and the mitochondrion. Thus, to determine the precise localization of HDP, we further modified the *hdp-mNG:loxP* parasites to episomally express minimal mCh-based reporters for both these organelles. No overlap was observed with the apicoplast marker, whereas HDP exhibited near-complete colocalization with the mitochondrial reporter (Figure 1F). HDP also coincided with the ΔΨ_mt_-dependent dye MitoTracker Red CMXRos (Figure 1F). Together, these results suggest that HDP is absent from the DV and instead localizes to the parasite mitochondrion throughout asexual blood stage development.

### HDP is critical for blood-stage survival

To assess the importance of HDP for the parasite’s blood-stage development, we treated *hdp-mNG:loxP* cultures with 20 nM rapamycin (RAP), triggering DiCre-mediated excision of the C-terminal two-thirds of the HDP coding sequence (Figure 1A). Diagnostic PCRs of the HDP locus confirmed RAP-induced loss of the target DNA compared to dimethyl sulfoxide (DMSO)-treated control parasites (Figure 2A). Loss of the mNG-tagged protein was verified by Western blot analysis. In DMSO-treated samples, mNG was detected in the detergent-resistant pellet fraction after parasite lysis, appearing as a band of ∼50 kDa, consistent with the expected size of the HDP-mNG fusion protein (52 kDa) (Figure 2B). An additional, more prominent band at ∼80 kDa was also observed, indicative of abnormal migration behavior during SDS-PAGE. Both bands were markedly depleted in lysates from RAP-treated parasites by the end of the first intraerythrocytic cycle, coinciding with the disappearance of mNG fluorescence in live microscopy, together confirming that HDP is efficiently removed upon RAP treatment (Figure 2B,C).

**Figure 2:**
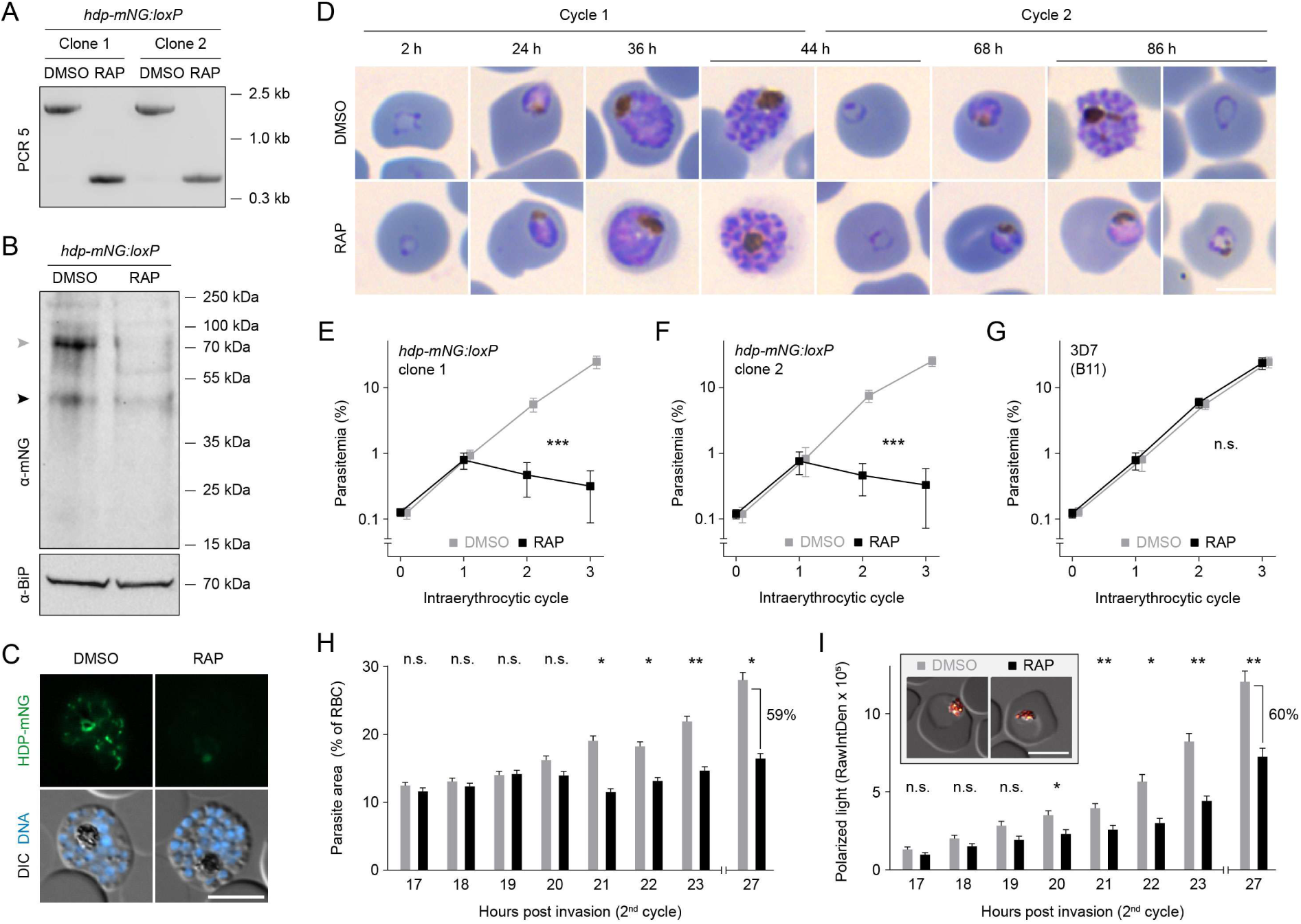
Loss of HDP causes developmental arrest accompanied by reduced hemozoin content. **(A-C)** RAP-induced loss of HDP, as confirmed by (A) diagnostic PCR5 (see Figure 1A), (B) Western blot, and (C) live cell imaging at the end of the first cycle. Black arrowhead: predicted size of HDP-mNG; grey arrowhead: higher molecular weight band. **(D)** HDP-deficient parasites arrest as trophozoites during the second cycle following RAP treatment, as shown by Giemsa staining. **(E-G)** HDP-deficient parasites fail to replicate beyond the second cycle. Growth curves are shown for two independent DMSO and RAP-treated *hdp-mNG:loxP* clones (E, F) and parental parasites (G). Shown are mean values +/- SD. n.s., non-significant; ***P<0.001, two-way ANOVA; N = 3 biological replicates with 3 technical replicates each. **(H, I)** Quantification of parasite size (H) and hemozoin content (I) demonstrates a tight temporal correlation between growth arrest and reduced hemozoin production in HDP-null parasites. Hemozoin was quantified by polarization microscopy. Shown are mean values +/- SEM. *P<0.05, **P<0.01, paired t-test; N = 99 parasites from 3 independent experiments. Inset: parasites 27 hours post-invasion, shown as a merge of DIC (grey) and polarized light (orange). Scale bars, 5 µm.

We next monitored parasite morphology in tightly synchronized cultures by light microscopy of Giemsa-stained blood smears over two intraerythrocytic cycles. While DMSO control parasites developed normally, RAP treated parasites completed the first cycle but arrested development in the second cycle as trophozoites (Figure 2D). Flow cytometry-assisted growth assays corroborated these observations, showing that RAP-treated parasites underwent one successful round of replication but failed to proliferate thereafter (Figure 2E,F). In contrast, the parental B11 line was unaffected by RAP treatment (Figure 2G). Combined, these findings indicate that HDP is essential for the survival of *P. falciparum* blood-stage parasites.

### Loss of hemozoin formation coincides with developmental arrest in HDP-null parasites

To more precisely determine when developmental arrest occurs following HDP knockout, we treated ring-stage cultures with DMSO or RAP, allowed them to develop into mature schizonts, and used these schizonts to initiate a new, highly synchronous invasion. Samples were then collected every two hours, and parasite size was quantified microscopically in methanol-fixed blood smears. RAP-treated parasites ceased normal growth at approximately 21 hours post-invasion in the second cycle (Figure 2H).

We analyzed the same parasite populations using polarization microscopy, which specifically visualizes birefringent hemozoin crystals and thereby provides a quantitative measure of hemozoin content. In DMSO-treated control parasites, hemozoin levels steadily increased over time, whereas RAP-treated parasites began producing less hemozoin at around 20 hours post-invasion, coinciding with the onset of developmental arrest (Figure 2I). At 27 hours post-invasion, RAP-treated parasites contained 60% of the hemozoin found in controls. With parasite size also reduced to 59%, HDP-deficient parasites maintained comparable hemozoin content relative to size, and the crystals remained normally distributed, accumulating in a single large DV (Figure 2H,I). The fact that a similar reduction in hemozoin content was well tolerated in an unrelated mutant (43) but coincides with death in HDP-deficient parasites further suggests that HDP has an essential function independent of heme biomineralization.

### HDP is critical for maintaining the mitochondrial membrane potential

In light of these findings, we hypothesized that loss of HDP would primarily affect mitochondrial physiology. To test this, we assessed ΔΨ_mt_ in DMSO and RAP-treated *hdp-mNG:loxP* parasites using MitoTracker Red CMXRos, which accumulates in mitochondria in a ΔΨ_mt_-dependent manner. As a positive control, stained parasites were treated with carbonyl cyanide-p-trifluoromethoxyphenylhydrazone (FCCP), an ionophore that collapses ΔΨ_mt_.

At 24 and 44 hours post-invasion in the first cycle, MitoTracker staining intensity was similar between DMSO- and RAP-treated parasites, while FCCP efficiently collapsed ΔΨ_mt_ under both conditions (Figure 3A). In the second cycle, at 12 hours post-invasion, RAP-treated parasites already displayed a slight, non-significant reduction in MitoTracker intensity. This reduction became more pronounced and statistically significant at 18 hours post-invasion and persisted at later time points, showing an approximately two-fold decrease relative to DMSO-treated parasites (Figure 3A,B). Importantly, the dissipation of ΔΨ_mt_ preceded the observed deficiencies in parasite size and hemozoin production (Figure 2H,I), indicating that mitochondrial depolarization upon loss of HDP is not a secondary consequence of parasite death.

**Figure 3:**
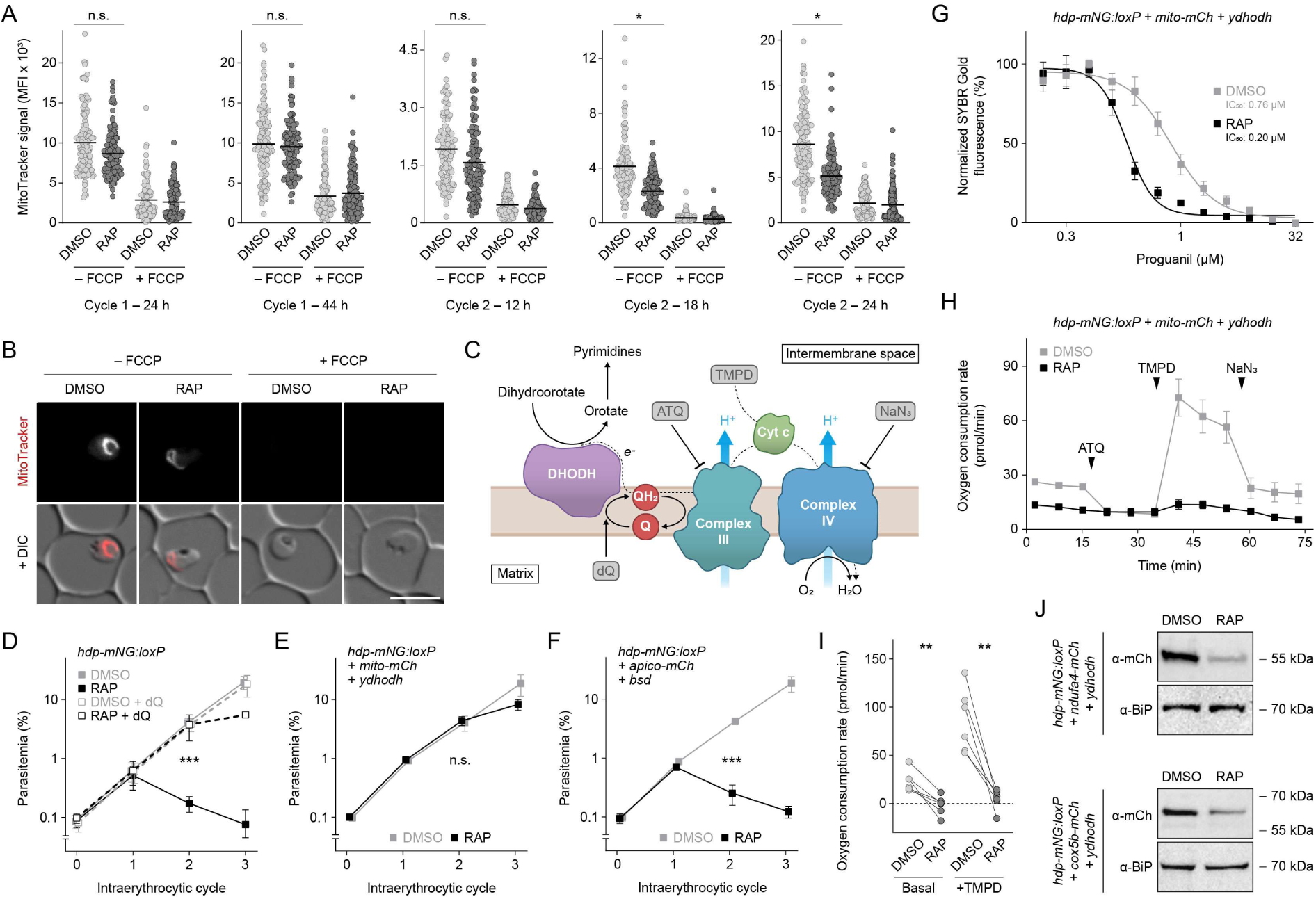
HDP is essential for mitochondrial electron transport. **(A,B)** Loss of HDP causes ΔΨ_mt_ dissipation. (A) Live microscopy of Mitotracker Red CMXRos staining was used to quantify ΔΨ_mt_ in the presence or absence of FCCP at the indicated timepoints. Mean and individual values are shown. n.s., non-significant; *P<0.05, paired t-test; N = 132 parasites from 4 independent experiments. (B) Representative microscopy images at 18 hours post-invasion in the second cycle. Scale bar, 5 µm. **(C)** Schematic representation of respiratory complexes III and IV and their role in pyrimidine biosynthesis. Exogenous compounds are shown in grey. DHODH, dihydroorotate dehydrogenase; Cyt c, cytochrome c; Q, ubiquinone; QH_2_, ubiquinol; dQ, decyl-ubiquinone; ATQ, atovaquone; TMPD, N,N,N′,N′-Tetramethyl-1,4-phenylendiamin; NaN_3_, sodium azide. **(D-F)** HDP-deficient parasites are rescued by exogenous dQ (D) or expression of yeast DHODH (yDHODH) (E), but not by expression of blasticidin S deaminase (BSD) (F). Growth curves show mean values +/- SD. ***P<0.001, two-way ANOVA; N = 3 biological replicates with 3 technical replicates each. **(G)** HDP-deficient parasites are hypersensitive to proguanil. Growth-response curves of yDHODH-rescued parasites are shown, with mean +/- SEM and IC_50_ values indicated. N = 3 biological replicates with 3 technical replicates each. **(H, I)** Loss of HDP disrupts respiration. (H) Oxygen consumption rate (OCR) of yDHODH-rescued parasites over the course of one Seahorse assay. Timepoints for compound addition are indicated. Shown are mean values +/- SEM. (I) OCR under basal conditions and in the presence of TMPD. Individual values are shown. **P<0.01, paired t-test; N = 6 assays with ≥ 6 technical replicates each. **(J)** HDP-deficient parasites show markedly reduced levels of complex IV subunits. Western blots of yDHODH-rescued *hdp-mNG:loxP* parasites expressing endogenous NDUFA4 (top) or Cox5B (bottom) fused to mCherry. Samples were harvested at the end of the first cycle.

### A metabolic respiratory chain bypass rescues HDP-deficient parasites

Our findings led us to hypothesize that HDP may play a role in the ETC, the main contributor to ΔΨ_mt_ (Figure 3C). Since the lethal effect of inhibiting mitochondrial respiration can be bypassed by alternative means of supporting dihydroorotate oxidation (10, 27), we tested whether exogenous supply of the electron acceptor decyl-ubiquinone (dQ) could rescue HDP-deficient parasites. Using standardized flow cytometry-assisted growth assays, we found that dQ enabled HDP-deficient parasites to replicate beyond the second cycle, indicating that restoration of DHODH activity was sufficient to compensate for the loss of HDP function (Figure 3D).

To further substantiate this, we tested whether expression of a ubiquinone-independent yeast dihydroorotate dehydrogenase (yDHODH) could similarly rescue HDP-null parasites. We thus analyzed replication of the *hdp-mNG:loxP + mito-mCh + ydhodh* mutant that was used for mitochondrial colocalization. This line expresses yDHODH as a drug-selectable marker, and we indeed found that it replicates efficiently even in the absence of HDP (Figure 3E). In contrast, *hdp-mNG:loxP + apico-mCh + bsd* parasites which express an unrelated blasticidin resistance cassette, still died in the second cycle after RAP treatment (Figure 3F). Together, these findings demonstrate that the loss of HDP can be largely compensated by bypassing the essential metabolic functions of the ETC.

### Severely reduced respiration upon loss of HDP

These phenotypic rescue experiments suggested a role for HDP in mitochondrial respiration. To explore this further, we tested whether HDP influences sensitivity to proguanil, an antimalarial known to synergize with ETC inhibitors and thought to target a complex III/IV-independent pathway that supports mitochondrial polarization (10). Using replication-based drug susceptibility profiling of the rescued *hdp-mNG:loxP + mito-mCh + ydhodh* line, we found that loss of HDP increased parasite sensitivity to proguanil by approximately four-fold (Figure 3G). This indicates that loss of HDP sensitizes the parasites to respiration-independent mitochondrial depolarization, as expected in case of impaired electron transport.

To directly assess respiratory function, we measured oxygen consumption in DMSO- and RAP-treated second-cycle *hdp-mNG:loxP + mito-mCh + ydhodh* parasites at young schizont stage, using an XFe96 flux analyzer. RAP-treated parasites displayed a markedly reduced oxygen consumption rate (OCR) compared to DMSO controls under basal conditions (Figure 3H,I). Addition of the complex III inhibitor atovaquone to DMSO-treated parasites lowered the OCR to the same basal level observed in RAP-treated samples, while having little additional effect on the RAP-treated parasites themselves, together indicating a near-complete block of respiration in the absence of HDP. Addition of N,N,N′,N′-tetramethyl-p-phenylenediamine dihydrochloride (TMPD), which bypasses complex III by donating electrons directly to cytochrome c, restored oxygen consumption in DMSO-treated parasites, raising it far above basal levels, but failed to do so in RAP-treated samples (Figure 3H,I). This suggests that the respiratory defect in HDP-deficient parasites arises, at least in part, from dysfunction downstream of complex III.

Suspecting that complex IV dysfunction might underlie the respiratory defect, we endogenously tagged the subunits NDUFA4 and Cox5B with mCh in a yDHODH-rescued *hdp-mNG:loxP* background. Western blotting revealed a substantial reduction of both subunits in HDP-deficient parasites already by the end of the first cycle, indicating widespread disorganization and loss of complex IV in the absence of HDP (Figure 3J).

### HDP is composed primarily of an adhesive protein domain

To better understand the role of HDP in the integrity and function of the ETC, we examined its molecular and structural features. HDP is predicted to consist primarily of a Fasciclin 1 (Fas1) domain, a widely conserved motif commonly involved in cell adhesion and, more rarely, in providing structural support within large protein assemblies (48, 49). Sequence alignment of HDP with representative Fas1-domain-containing proteins across diverse taxa revealed several conserved residues, including the histidine of the YH motif and residues within the H1 and H2 regions (48) (Figure 4A).

**Figure 4:**
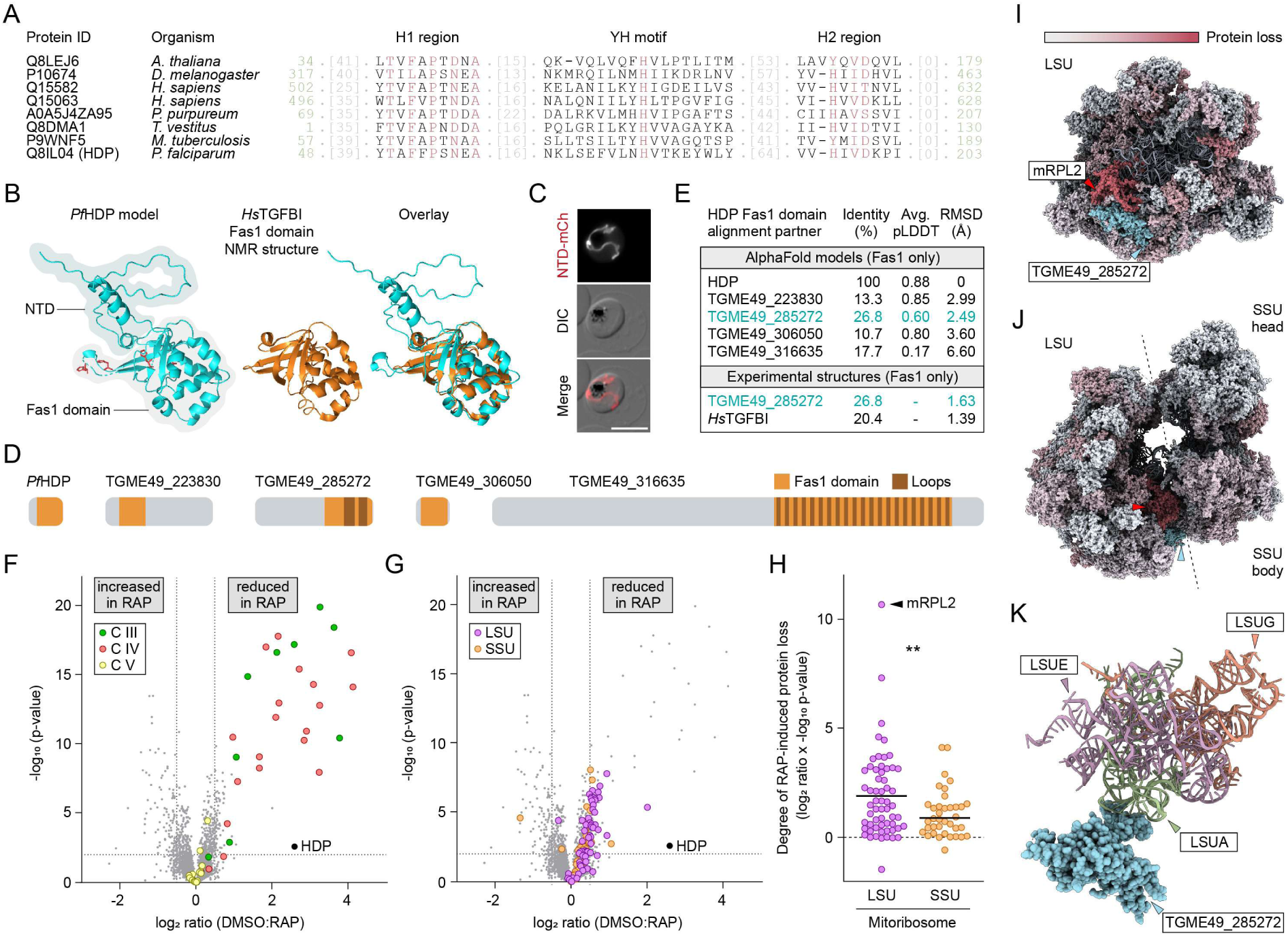
HDP participates in mitochondrial protein biosynthesis. **(A)** Multiple sequence alignment of HDP with Fas1 domain-containing proteins from other organisms. Semi-conserved regions H1 and H2 and the YH motif are shown. Red residues indicate high conservation. Grey numbers mark omitted regions, green numbers denote Fas1 domain boundaries. **(B)** HDP consists of a conserved Fas1 domain and a largely unstructured N-terminal domain (NTD). The AlphaFold model of HDP was aligned to the NMR structure of the human Transforming Growth Factor Beta Induced (TGFBI) Fas1 domain. Histidine residues previously implicated in β-hematin formation are highlighted in red. **(C)** The NTD of HDP is sufficient for targeting mCherry to the mitochondrion. Scale bar, 5 µm. **(D)** Domain organization of HDP compared with Fas1 proteins from *Toxoplasma gondii*. **(E)** Sequence and structural alignments of HDP with *T. gondii* Fas1 proteins identify TGME49_285272 as the most likely HDP homolog. Sequence identity, average AlphaFold pLDDT scores, and RMSD values from structural alignments are shown. **(F-H)** Whole-cell proteomics of yDHODH-rescued *hdp-mNG:loxP* parasites. (F) Subunits of complexes III (C III) and IV (C IV) are severely reduced in HDP-deficient parasites, whereas complex V (C V) remains unaffected. (G) Components of the mitoribosome’s large (LSU) and small (SSU) subunits are destabilized upon loss of HDP. Note that (F) and (G) show the same dataset with different proteins highlighted. Dashed lines indicate significance and fold-change cut-offs. Benjamini-Hochberg corrected t-test; N = 4 independent biological replicates with 3 technical replicates each. (H) Loss of HDP affects the LSU more strongly than the SSU. Mean and individual values are shown for a combined score incorporating fold-change and significance. **P<0.01, unpaired t-test; N = 59 LSU, 38 SSU proteins. **(I-K)** HDP-dependent reduction of *P. falciparum* mitoribosomal proteins mapped onto the *T. gondii* mitoribosome structure (22). Proteins are color-coded to reflect the reduction of their respective *P. falciparum* orthologs following HDP knockout. (I) The LSU shown from the inter-subunit side. (J) The assembled mitoribosome with LSU and SSU in contact. The putative HDP ortholog TGME49_285272 is highlighted in cyan. (K) TGME49_285272 contacts ribosomal RNAs. Only TGME49_285272 and LSU rRNAs are shown, with the rest of the *T. gondii* mitoribosome omitted for clarity.

Structural prediction of HDP using AlphaFold 3 (50) produced a high-confidence model for the Fas1 domain, which aligned closely with the experimentally determined structure of a human Fas1 domain (51) (Figure 4B). In addition, HDP possesses a 48-residue N-terminal domain (NTD) enriched in positively charged and hydrophobic residues, consistent with a mitochondrial targeting signal. The NTD was modeled with low confidence and lacked a well-defined fold (Figure 4B). When fused to mCh, these 48 residues directed the reporter to the mitochondrion, supporting a role for the HDP N-terminus in mitochondrial targeting (Figure 4C).

HDP is the only protein encoded in the *P. falciparum* genome predicted to contain a Fas1 domain. In contrast, the related coccidian parasite *Toxoplasma gondii* encodes four such proteins: one of comparable size and overall architecture, TGME49_306050, and three significantly larger proteins (Figure 4D). Among the latter, TGME49_285272 was recently identified as a component of the mitoribosome large subunit (LSU) (21, 22). However, it contains an NTD more than eight times longer than that of HDP, and its Fas1 domain includes two extended loops absent from the *Plasmodium* protein (Figure 4D, Supplementary Figure S1). Despite these differences in overall architecture, the Fas1 domain of TGME49_285272 exhibited the highest sequence similarity to that of HDP among all *T. gondii* Fas1 proteins (Figure 4E). The AlphaFold model of TGME49_285272 Fas1 also showed the greatest structural overlap with that of HDP, whereas the remaining *T. gondii* proteins aligned less closely – though this may partly reflect differences in model confidence (Figure 4E, Supplementary Figure S1). Finally, comparison with the experimentally determined structure of the TGME49_285272 Fas1 domain yielded the closest match to HDP, with only the human Fas1 domain achieving slightly better alignment (Figure 4E, Supplementary Figure S1).

### HDP participates in mitochondrial protein biosynthesis

With the mitoribosomal protein TGME49_285272 identified as the most likely HDP ortholog, we predicted that loss of HDP would severely impair mitochondrial protein biosynthesis, the process responsible for producing the complex III component cytochrome b and the complex IV components Cox1 and Cox3 (19, 20). Consequently, HDP loss would be expected to destabilize not only complex IV, as reflected by reduced NDUFA4 and Cox5B (Figure 3J), but also complex III. To test this, we performed whole-cell proteome analysis of DMSO- versus RAP-treated *hdp-mNG:loxP + ydhodh* parasites. Nearly all subunits of complexes III and IV were markedly reduced upon HDP inactivation, with Cox1 and the complex IV component C4AP5 being completely undetectable in RAP-treated parasites (Figure 4F, Supplementary Table S2). In contrast, complex V, whose components are nuclear-encoded and imported into the mitochondrion post-translationally, remained unaffected, supporting a specific role of HDP in mitochondrial protein synthesis.

Among the proteins significantly reduced in HDP-deficient parasites were also multiple mitoribosomal components. Indeed, most mitoribosomal proteins showed lower abundance, though not all reached statistical significance (Figure 4G, Supplementary Table S2). This was generally more pronounced in the LSU than the SSU, with the LSU protein mRPL2 (PF3D7_1132700) being most affected (Figure 4H). Mapping these proteins onto the recently determined structure of the *T. gondii* mitoribosome revealed that the ortholog of mRPL2 is in direct contact with TGME49_285272, the protein we identified as the most likely HDP ortholog (Figure 4I). Both proteins localize to the LSU–SSU interface, potentially explaining why loss of HDP affects both subunits (Figure 4I-K). Destabilization was widespread across the LSU and SSU, indicating broad mitoribosomal instability upon HDP inactivation (Figure 4I,J; Supplementary Figure S2).

Collectively, these results suggest that HDP is an integral component of the LSU and critical for mitochondrial protein biosynthesis, which produces essential components for the stability and function of the malaria parasite’s ETC.

### HDP does not function in heme biomineralization

Finally, we revisited the previously proposed role of HDP in heme biomineralization by analyzing the quantity and morphology of hemozoin in metabolically rescued *hdp-mNG:loxP + mito-mCh + ydhodh* parasites. This approach allowed us to re-examine the pathway without confounding effects from developmental retardation. Parasites were analyzed 36 hours post-invasion in the second cycle, a time point at which HDP had been depleted for some time (Figure 2B,C), and well beyond when parasites lacking yDHODH would have died (Figure 2D,H). Quantitative polarization microscopy revealed no differences in either the localization or abundance of hemozoin between DMSO- and RAP-treated parasites (Figure 5A-C). Consistent with this, scanning electron microscopy of isolated hemozoin showed no HDP-dependent changes in crystal size or morphology, with the crystals maintaining their characteristic brick-like shape (Figure 5D). Moreover, parasite sensitivity to chloroquine, an established heme biomineralization inhibitor, remained unchanged in the absence of HDP (Figure 5E).

**Figure 5:**
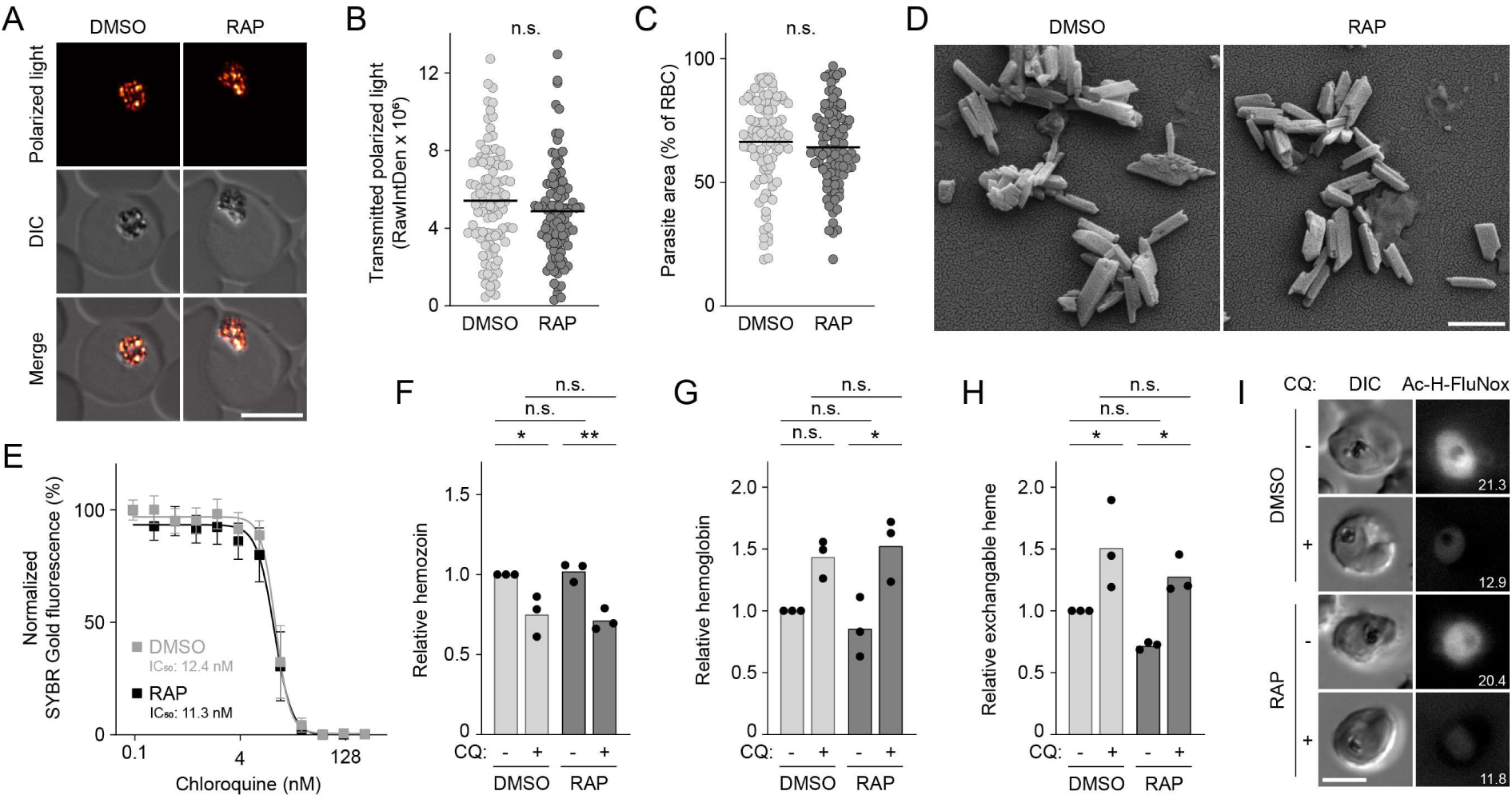
Loss of HDP does not affect hemozoin formation in yDHODH-rescued parasites. To assess heme biomineralization independent of developmental arrest, metabolically rescued parasites were analyzed in the second cycle post-induction, using the *hdp-mNG:loxP + mito-mCh + ydhodh* line (A-E) or the alternative *hdp-smHA:loxP + ydhodh* NF54 line (F-I). **(A-C)** Loss of HDP does not alter hemozoin quantity, as determined by polarization microscopy (A, B), or parasite size (C). Mean and individual values are shown. n.s., non-significant; paired t-test; N = 99 parasites from 3 independent experiments. Scale bar, 5 µm. **(D)** Hemozoin morphology is unchanged in the absence of HDP, as shown by scanning electron microscopy of purified crystals. Scale bar, 500 nm. **(E)** HDP-deficient parasites exhibit normal chloroquine sensitivity. Growth-response curves are shown with mean +/- SEM and IC_50_ values indicated. N = 3 biological replicates with 3 technical replicates each. **(F-I)** HDP deletion does not alter the distribution of different heme species. Heme associated with hemozoin (F), hemoglobin (G), or the exchangeable heme pool (H) was quantified using a pyridine-based heme fractionation assay. Chloroquine (CQ) was used as a positive control. Details on drug addition and analysis time points are provided in the Methods section. Mean and individual values are shown. *P<0.01; **P<0.01; N = 3 biological replicates with 2 technical replicates each. (I) Ac-H-FluNox staining reveals no effect of HDP deletion on cytoplasmic free-heme levels. Numbers indicate mean intensity values. N ≥ 136 parasite from 5 independent experiments (see Supplementary Figure S4). Scale bar, 5 µm.

Because the N-terminal 72 amino acids of HDP remain intact in our 3D7-based knockout, we next sought to rule out a potential contribution of these residues to hemozoin formation. To this end, we generated an independent DiCre line based on the alternative *P. falciparum* strain NF54, flanking the entire HDP coding sequence with *loxP* sites and tagging the C-terminus with a spaghetti monster haemagglutinin tag (smHA) (Supplementary Figure S3A). A yDHODH expression cassette was again introduced to bypass HDP-dependent respiratory defects. Correct genome editing and RAP-induced HDP loss were confirmed by diagnostic PCR, Oxford Nanopore long-read sequencing, whole-genome sequencing, and Western blot (Supplementary Figure S3B-D). The resulting *hdp-smHA:loxP + ydhodh* NF54 line replicated efficiently following HDP inactivation and recapitulated the proguanil hypersensitivity phenotype observed in the 3D7-based mutant (Supplementary Figure S3E,F).

To obtain detailed profiles of heme distribution, we performed a pyridine-based heme fractionation assay with DMSO and RAP-treated *hdp-smHA:loxP + ydhodh* NF54 parasites, quantifying heme associated with hemozoin, hemoglobin, and the residual exchangeable heme pool, with chloroquine treatment serving as a positive control (52). As expected, chloroquine affected all heme species; however, the loss of HDP had no effect on any of the heme pools and did not alter the magnitude of chloroquine-induced perturbation (Figure 5F-H). We further complemented this analysis with live-cell imaging of Ac-H-FluNox, a cell-permeable fluorescent probe specific for labile heme (53). Although chloroquine treatment quenched Ac-H-FluNox fluorescence, as previously reported (54), the signal remained unaffected by HDP inactivation (Figure 5I, Supplementary Figure S4). Consistent with our findings in the 3D7-based line, excision of the entire HDP gene did not alter chloroquine sensitivity in the *hdp-smHA:loxP + ydhodh* NF54 parasites (Supplementary Figure S3G).

Collectively, our findings indicate that, despite previous reports of HDP-catalyzed β-hematin formation *in vitro*, HDP does not contribute to hemozoin production in the parasite across different *P. falciparum* genetic backgrounds. Instead, our in-depth analysis of mitochondrial physiology places the protein in the mitoribosome.

## DISCUSSION

Since Robert Virchow and Charles Laveran first recognized the connection between hemozoin and malaria (55, 56), generations of parasitologists have sought to understand how *Plasmodium* parasites orchestrate the formation of such highly ordered crystalline structures – a process that later turned out to be the target of several highly effective antimalarials (33, 35, 36). Early clues that parasite proteins might contribute to hemozoin formation came from observations that parasite lysates could catalyze β-hematin formation *in vitro* (57). In the search for specific parasite proteins capable of mediating this process, the first candidates examined were HRPII and III (41). Both proteins are present in the DV and contain repetitive histidine-rich stretches resembling heme-binding motifs. Although both proteins can efficiently catalyze β-hematin formation *in vitro*, they are dispensable for parasite survival, and their absence does not affect hemozoin production *in cellulo* (41). Recent evidence indicates that HRPII instead assembles into heme-rich nanoparticles, which are released into the bloodstream and trigger inflammatory responses in endothelial cells (58).

The subsequent discovery of HDP as a potent inducer of β-hematin formation provided an alternative candidate for this essential parasite pathway (42). Although no clear rationale has ever been given for examining this protein in this context, detailed biochemical studies have since aimed to elucidate the underlying reaction chemistry, resulting in a model in which specific histidine residues of HDP orient two hematin monomers to promote their dimerization (59–62).

Early localization studies using immunofluorescence and immuno-electron microscopy detected HDP in the host cell cytoplasm, hemoglobin transport vesicles, and the DV, consistent with a trafficking pathway involving secretion, export, and reuptake (42). When expressed episomally as a C-terminal GFP fusion, HDP localized to a tubular compartment within the parasite, interpreted as a hemoglobin trafficking intermediate (45). Support for a role in the DV came from Co-IP experiments showing that HDP associates with hemoglobin-degrading proteases, suggesting that they may act in concert with HDP to coordinate heme release and sequestration (44).

In our study, we tagged the endogenous HDP locus and found the protein to be associated with the parasite mitochondrion, colocalizing with both MitoTracker and a well-established mitochondrial reporter (63, 64). Low-level signals observed in the DV corresponded to natural background fluorescence, and even strong overexpression did not reveal any secondary protein pool. Since HDP is essential for parasite survival, the ability to tag the endogenous protein without compromising viability further supports that the observed localization reflects the physiological distribution of HDP. We found that the NTD of HDP is sufficient for mitochondrial targeting, whereas the N-termini of DV proteins typically direct them into the secretory pathway (65, 66). The most compelling support for HDP functioning exclusively in the mitochondrion comes from our observation that metabolically bypassing respiratory dysfunction fully restores the development of HDP-deficient parasites, along with normal hemozoin production.

The predicted structure of HDP indicates that two of the four histidine residues (H122 and H197), previously proposed to coordinate hematin binding (59), are deeply buried within the Fas1 fold, suggesting they are not accessible in solution (Figure 4B). Preparations of HDP retain their activity even after heating to >90°C, indicating that the protein’s tertiary structure may not be critical for catalysis (42, 61). Circular dichroism spectroscopy, however, revealed some thermostability (42), in line with our observation of abnormal SDS-PAGE migration, which may reflect an inability to fully denature the tightly folded Fas1 domain. The catalytic role of HDP *in vitro* likely reflects the promiscuous nature of β-hematin formation, which can be greatly facilitated by nucleation centers supplied by proteins such as HDP and the HRPs, as well as by buffer components, impurities, certain lipids or ordered surfaces. Supporting this notion, studies in mice showed that loss of a lysosomal heme exporter leads to hemozoin deposits in reticuloendothelial macrophages that digest aged RBCs (67). Heme accumulation in acidic lysosomes is therefore sufficient to trigger biomineralization, even in organisms that do not normally form hemozoin, indicating that crystallization is chiefly driven by heme supersaturation and pH, with proteins, lipids, and small molecules only modulating the process. With HDP effectively disqualified, the only *Plasmodium* protein currently known to affect hemozoin production *in cellulo* is PV5, a DV-localized lipocalin. Its inactivation reduces hemozoin formation by approximately two-fold and causes excessive branching of the crystals, with only mild effects on parasite viability (43).

Our comprehensive analysis of mitochondrial physiology demonstrates a role of HDP in mitochondrial protein biosynthesis and mirrors previous observations made upon inactivation of the LSU protein mRPL13 (68). Loss of HDP destabilizes the mitoribosome, thereby preventing the production of key components of complexes III and IV. As a result, electron transport through the ETC is completely blocked, dissipating the pH gradient required for ΔΨ_mt_ maintenance and sensitizing the parasite to further mitochondrial depolarization. Several lines of evidence support the notion that HDP is an integral part of the mitoribosomal LSU: (1) despite notable differences in protein size and architecture, structural and sequence alignments identify HDP as the most likely ortholog of the *T. gondii* LSU protein TGME49_285272; (2) HDP inactivation selectively affects complexes whose integrity depends on mitochondrial protein synthesis (19, 20); and (3) HDP inactivation compromises mitoribosome stability, particularly within the LSU, where a predicted TGME49_285272 neighbor protein is most affected.

A possible, though less likely, function for HDP could be the direct stabilization of complexes III and IV or of their shared supercomplexes. Indeed, complexome profiling has revealed the presence of respiratory supercomplexes in *P. falciparum* gametocytes, although their existence in asexual blood stages is less clear (69). In *T. gondii*, supercomplexes of complexes III and IV have been observed, but disrupting the underlying interactions caused only mild defects in parasite fitness and did not affect electron flow through the ETC (70). In any case, a direct role for HDP in stabilizing the ETC would be difficult to reconcile with the observed mitoribosomal destabilization.

In *T. gondii*, TGME49_285272 is located at the interface side of the LSU (21, 22). Available structures of the mitoribosome resolved only 227 of the protein’s 706 residues, covering the Fas1 core but excluding the two loops emerging from the domain and lacking most of the extended NTD. Interestingly, the resolved NTD residues are conserved in *P. falciparum*, spanning the entire NTD of HDP except for the first eleven residues. This suggests that, despite holding targeting-relevant information, the majority of the NTD remains intact following mitochondrial import. This is consistent with observations in yeast, where approximately 25% of mitoribosomal proteins possess internal targeting motifs that are not cleaved but are rich in positively charged and hydrophobic residues to facilitate import (71). A stretch of the TGME49_285272 NTD corresponding to HDP residues 22-31 appears to align closely with the rRNA LSUA in the *T. gondii* mitoribosome structure, suggesting a potential role in rRNA stabilization (21, 22) (Figure 4K). The exact function of the Fas1 domain remains unclear, but it likely contributes to stabilizing the overall LSU structure through its adhesive properties, consistent with the widespread destabilization observed upon HDP loss. Indeed, a similar role has been shown for Fas1-domain-containing linker proteins that stabilize light-harvesting complexes in algae (49).

Surprisingly, loss of HDP caused parasites to arrest development only in the second cycle, even though complex IV subunits were already strongly reduced by the end of the first cycle. At that point, ΔΨ_mt_ remained intact, suggesting that despite the impact of HDP loss on the ETC, electron transport was still sufficient to maintain ΔΨ_mt_, or that processes dissipating the proton gradient had not yet taken effect. Moreover, pyrimidine levels produced before mitochondrial dysfunction may have been enough to sustain the parasites through one intraerythrocytic cycle, even after ubiquinone depletion caused pyrimidine synthesis to halt. Parasites ultimately died at the trophozoite stage, when pyrimidine demand reaches a critical level for the first round of DNA replication.

Together with its essential role in asexual blood-stage development, the divergent composition and structure of the apicomplexan mitoribosome highlights this unique protein-RNA assembly as an attractive target for future drug development (21, 22, 68, 72). Current antimalarial strategies already exploit selective inhibition of translation within the parasite’s endosymbiotic organelles, with compounds such as doxycycline, clindamycin, and azithromycin targeting the apicoplast ribosomes (73–75). Secondary effects on mitochondrial translation have been observed, especially at higher concentrations (76–78), underscoring the mitoribosome’s potential as a largely unexplored drug target. Elucidating the *Plasmodium* mitoribosome structure, as previously achieved for *T. gondii*, could pave the way for the rational design of inhibitors targeting the parasite’s mitochondrial translation machinery. Such an approach would provide a powerful new avenue for antimalarial combination therapies that exploit synergy with established drugs acting on respiration, ΔΨ_mt_ maintenance, or pyrimidine biosynthesis (72).

## METHODS

### Parasite cultivation and synchronization

*P. falciparum* blood-stage parasites were cultured in deidentified human RBCs (B+; Universitätsklinikum Hamburg-Eppendorf) at 1-5% hematocrit in Roswell Park Memorial Institute medium 1640 (RPMI 1640) supplemented with 0.5% AlbuMAXII (Thermo Fisher Scientific) at 37°C under 1% O_2_, 5% CO_2_, and 94% N_2_. For synchronization, mature schizonts were enriched by Percoll density gradient centrifugation and added to fresh RBCs for a 2-hour invasion under constant agitation. Residual schizonts were removed with a second Percoll gradient followed by 5% sorbitol treatment. For reverse genetic experiments, synchronized cultures were treated immediately with 20 nM RAP (Clontech Laboratories) or DMSO (Merck).

### Generation and validation of transgenic parasites

For generation of the 3D7-based *hdp-mNG:loxP* mutant, a Cas9 repair template was generated by commercial gene synthesis, and the mNG tag was cloned in-frame using HpaI and EcoRV restriction sites. DiCre-expressing *P. falciparum* schizonts (strain B11) (47) were transfected with 20 µg of guide plasmid to induce Cas9-mediated double-strand cleavage, together with 60 µg of linearized repair template, using a 4D Nucleofector system (Lonza). Transgenic parasites were selected with 10 nM WR99210 (Jacobus Pharmaceuticals).

For the *hdp-smHA:loxP + ydhodh* NF54 line, the repair template was generated by assembly PCR and introduced into XhoI/EagI-digested yPM2GT vector (79) by In-Fusion cloning. The repair template, together with the guide plasmid, was transfected into the NF54^attB-DiCre^ line (80). Transgenic parasites were selected with 0.9 µM DSM1 (Merck). The conditional knockout mutants were isolated by limiting dilution and validated using multiple complementary approaches, including diagnostic PCR, Oxford Nanopore long-read sequencing, whole-genome sequencing, Western blotting, and fluorescence microscopy, with each mutant confirmed by a subset of these methods.

Endogenous tagging of Cox5B (PF3D7_0927800) and NDUFA4 (PF3D7_1439600) was achieved using the selection-linked integration system (81). Tagging vectors were derived from the pSLI-PF3D7_1476700-mNeonGreen-glmS plasmid (82) and generated by first replacing mNG with mCh using the MluI and SalI sites, followed by cloning of the respective C-terminal region in frame with mCh using the NotI and MluI sites. After transfection, parasites were selected with WR99210 until resistant populations emerged and then treated with 0.4 mM neomycin (Merck) to enrich for integrants. Plasmids for episomal expression of the apicoplast (83) and mitochondrial (64) markers were maintained using 2 µg/mL blasticidin S (Thermo Fisher Scientific) or 0.9 µM DSM1, respectively. The episomal construct driving NTD-mCh expression was generated by replacing the citrate synthase leader sequence in the mitochondrial marker plasmid pNMD3:Mito-mCherry-DHODH (64) with the first 48 residues of HDP, using the XhoI and KpnI sites.

All transgenic parasite lines generated in this study are listed in Supplementary Table S3, and primers for molecular cloning, guide plasmid construction, diagnostic PCR, and the repair template sequences are provided in Supplementary Table S2.

### Western blotting

For Western blot analysis of 3D7-based mutants, parasites were released from RBCs by treatment with 0.03% saponin in PBS, pelleted, and lysed in 0.5 x PBS containing 4% SDS, 0.5% Triton X-100, and protease inhibitor cocktail (Roche). After one freeze-thaw cycle, lysates were centrifuged at 21,130 x g for 10 minutes. Supernatant and pellet fractions were taken up in 5x Laemmli buffer containing 0.6 M dithiothreitol, heated to 85°C for 5 minutes, and subjected to SDS-PAGE. Proteins were transferred onto nitrocellulose membranes, blocked with milk, and probed with α-BiP (rabbit, 1:2,000) (84), α-mNG (ChromoTek, mouse, 1:1,000) or α-RFP (ChromoTek, rat, 1:1,000) primary antibodies. Horseradish-peroxidase-conjugated secondary antibodies (Agilent, 1:5,000 - 1:3,000) and ECL reagent were used for chemiluminescence detection. Western blot analysis of *hdp-smHA:loxP + ydhodh* NF54 parasites was performed as previously described (54), using α-HA (Merck, rabbit, 1:1,000) and α-aldolase (abcam, rabbit, 1:1,000) primary antibodies.

### Parasite replication and drug sensitivity assays

As described previously (85), growth assays were initiated with synchronized ring-stage cultures adjusted to a parasitemia of 0.1% and supplemented with DMSO, RAP and/or 25 µM dQ (Cayman Chemical). Parasitemia was quantified every two days over four replication cycles by flow cytometry. For this, parasites were stained for 40 minutes at 37°C in medium containing SYBR Green nuclear dye (1:10,000; Thermo Fisher Scientific), followed by a 40-minute de-staining period. The proportion of stained cells was then measured on an automated NovoCyte 3000VYB flow cytometer equipped with a NovoSampler Pro (Agilent).

To analyze drug sensitivity of the *hdp-mNG:loxP + mito-mCh+ ydhodh* mutant, synchronized parasite cultures were adjusted to 0.1% parasitemia at 2% hematocrit and incubated in 96-well black-bottom plates (Thermo Fisher Scientific) with DMSO or RAP and varying concentrations of chloroquine or proguanil (both from Merck) in a final volume of 200 µl. Parasites were lysed 120 hours post-invasion, and DNA was stained with SYBR Gold (Invitrogen), as described previously (86). Fluorescence was measured using a GENios plate reader (Tecan), with excitation and emission wavelengths set to 485 nm and 535 nm, respectively. Raw fluorescence values were corrected using non-infected RBC controls and normalized to the highest and lowest readings.

Dose-response assays with *hdp-smHA:loxP + ydhodh* NF54 parasites were performed as previously described, with minor modifications (87). Parasites pre-treated with DMSO or RAP were seeded at 0.3% parasitemia and 1% hematocrit in 96-well flat-bottom plates and exposed to serial drug dilutions for 72 hours. Plates were subsequently analyzed by flow cytometry as previously described (54).

### Microscopy

All light and fluorescence microscopy was performed on a DM6 B microscope equipped with a K8 B/W camera and a K3C color camera (all from Leica), unless stated otherwise. The microscope was fitted with two crossed polarizers to enable selective visualization of hemozoin. For quantification of parasite size and hemozoin content, methanol-fixed blood smears were recorded in the DIC and polarization channels. RBC and parasite were manually outlined, and their areas measured. Hemozoin clusters were delineated in the polarization channel using manual thresholding, and their raw integrated densities were measured and corrected for background.

To assess ΔΨ_mt_, cells were stained at 37°C for 30 minutes in medium containing 2.5 nM MitoTracker Red CMXRos (Invitrogen), with or without 250 µM FCCP (Merck). After 30 additional minutes of de-staining, parasites were then imaged, and mitochondria were outlined using manual thresholding to determine background-corrected mean fluorescence intensity. Vacuolar autofluorescence was quantified by measuring background-corrected raw integrated density in manually outlined DVs, excluding parasites in which the mitochondrion was in close proximity to the DV.

To examine crystal morphology, hemozoin was isolated from 36-hour-old *hdp-mNG:loxP + ydhodh* parasites in the second cycle post-induction, as described previously (43). Parasites were lysed in 2% SDS, and the lysate was centrifuged at 21,130 x g for 5 minutes. The resulting pellet was washed twice in 2% SDS and three times with distilled water. Crystals were transferred onto 12 mm round glass coverslips and allowed to dry. Coverslips were then mounted on specimen stubs, sputter-coated with 5 nm gold using a CCU-010 compact coating unit (safematic), and then imaged using a CLARA scanning electron microscope (Tescan) at 5 keV.

For the analysis of free heme *in cellulo*, *hdp-smHA:loxP + ydhodh* NF54 parasites were treated with DMSO or RAP at the ring stage and then incubated with or without 150 nM chloroquine 72 hours post-induction for 5 hours. Parasites were subsequently stained with 10 µM Ac-H-FluNox for 30 minutes at 37°C, as previously described (54). Cells were washed twice with PBS and imaged using an Imager M2 Plus widefield fluorescence microscope (Zeiss) at excitation and emission wavelengths of 470 and 509 nm, respectively. Parasites were manually outlined, and background-corrected mean fluorescence intensity was quantified.

### Extracellular flux analysis

For Seahorse assays, 30-mL cultures containing young schizonts at >3% parasitemia and 4% hematocrit were collected in the second cycle post-induction. Parasites were released from their host cells by incubation in PBS containing 0.05-0.15% saponin (Merck). The isolated parasites were washed twice in PBS and resuspended in Mitochondria Assay Solution (MAS; 220 mM mannitol, 70 mM sucrose, 10 mM KH_2_PO_4_, 5 mM MgCl_2_, 2 mM HEPES, and 1 mM EGTA, adjusted to pH 7.2), supplemented with 2 mg/mL fatty acid-free bovine serum albumin, 24 mM malate, and 0.1% DMSO (all from Merck). Parasites were counted using a Neubauer chamber and resuspended at 5 x 10^7^ cells per mL in supplemented MAS further containing 0.002% digitonin (Merck), as described previously (64, 88).

OCR was measured on a Seahorse XFe96 Analyzer using Wave software (Agilent). For each well, 5 x 10^6^ parasites were loaded into CellTak-coated plates (Corning) and allowed to adhere by gentle centrifugation at 800 x g for 10 minutes. Even distribution and equal parasite density were confirmed microscopically. The assay included a total of 12 measurement cycles, each consisting of 3 minutes of mixing followed by 3 minutes of measuring. Three cycles were recorded sequentially for each step: first to assess basal respiration, then following the injection of 5 µM atovaquone, next after the addition of 0.2 mM TMPD with 2 mM ascorbate, and finally after 10 mM sodium azide (all from Merck). Assays were repeated with 6 independent parasite preparations, each comprising 6-30 technical replicates per condition, depending on cell yield.

### Label-free quantitative mass spectrometry

For whole-proteome profiling, 50 mL of DMSO- and RAP-treated *hdp-mNG:loxP + ydhodh* ring-stage cultures were adjusted to a parasitemia of 1.5% and a hematocrit of 2%, then harvested 80 hours post-invasion. Parasites were released from RBCs by a 10-minute incubation in ice-cold PBS containing 0.03% saponin, pelleted, and washed twice in the same buffer. Freed parasites were washed three additional times in ice-cold PBS. After a final centrifugation, the supernatant was removed and the parasite pellets were snap frozen in liquid N_2_ and subsequently stored at -70°C. Parasite proteins were isolated, digested with trypsin, and analyzed by liquid chromatography – mass spectrometry (LC-MS), as described in the Supplementary Methods.

### Heme fractionation assay

For the heme fractionation assays, *hdp-smHA:loxP + ydhodh* NF54 parasites induced at the ring stage were incubated with or without 100 nM chloroquine 56 hours post-induction for 16 hours. Parasites were released from RBCs with 0.035% saponin, washed three times with PBS to remove host cell hemoglobin, and resuspended in 100 µL PBS. Parasite concentration was determined by flow cytometry using a 5 µL aliquot stained with 1:25,000 Acridine Orange and supplemented with 25 µL CountBright Absolute Counting Beads (both from Thermo Fisher Scientific). Samples were analyzed on an Attune NxT Acoustic Focusing Cytometer (Thermo Fisher Scientific), and parasite numbers were calculated according to the manufacturer’s instructions.

Heme species were quantified using the remaining 95 µL of the cell suspension, as previously described (52). Briefly, cells were hypotonically lysed by addition of 50 μL water, sonicated, and mixed with 50 μL 0.2 M HEPES (pH 7.5; Merck) and 50 µL water. Samples were centrifuged at 4,500 x g for 20 minutes, and the supernatant corresponding to the hemoglobin fraction was supplemented with 50 μL 4% SDS (Merck), sonicated, incubated at 37°C for 30 minutes, and then mixed with 50 μL 0.3 M NaCl and 50 μL 25% pyridine (Merck). The pellet was resuspended in 50 μL water and 50 μL 4% SDS, sonicated, and incubated at 37°C for 30 minutes to solubilize the exchangeable heme. Following addition of 50 μL 0.2 M HEPES, 50 μL 0.3 M NaCl, and 50 μL 25% pyridine, samples were centrifuged and the supernatant was collected as the exchangeable heme fraction. The remaining pellet was resuspended in 50 μL water and 50 μL 0.3 M NaOH, sonicated, and incubated at 37°C for 30 minutes to solubilize hemozoin. Then, 50 μL 0.2 M HEPES, 50 μL 0.3 M HCl, and 50 μL 25% pyridine were added along with 150 μL of water.

Heme species were quantified using a standard curve prepared from bovine hematin (Merck) in 0.3 M NaOH. Absorbance by the heme-pyridine complex was recorded at a wavelength of 405 nm on an EnVision Multimode Plate Reader (PerkinElmer). Heme content was calculated by dividing the total heme in each sample by the parasite number, and values were normalized to the DMSO control.

### Statistical analysis and software

Statistical analyses were performed using GraphPad Prism (version 10.4.1). Comparisons between DMSO- and RAP-treated conditions were evaluated using paired t-tests, while unpaired t-tests were applied for comparisons between two independent groups. Multiple group comparisons were analyzed by one-way ANOVA with Tukey’s post hoc test, and growth curves were assessed by two-way ANOVA. For imaging-based experiments, statistical tests were applied to the mean values of independent biological replicates rather than to individual parasites. IC_50_ values were calculated by non-linear regression using the “log(inhibitor) vs. response – Variable slope (four parameters)” model.

Raw mass spectrometry data were processed using DIA-NN (version 2.2.0, see Supplementary Methods) (87). Proteins were included in the final datasets only if identified by at least two peptides. Statistical analysis was performed using Student’s t-test with Benjamini-Hochberg correction, controlling the false discovery rate at 1%. Proteins were considered differentially expressed if they showed a log_2_ fold-change ≥ 0.5 or ≤ -0.5, an adjusted p-value ≤ 0.01, and were detected in more than 75% of runs in at least one condition. Combined reduction scores were calculated as the log_2_ fold-change multiplied by the negative log_10_ of the p-value.

Microscopy images were analyzed and processed using FIJI (version 2.16.0/1.54p). Structural models of HDP and Fas1-domain-containing proteins of *T. gondii*, as well as individual Fas1 domains, were generated using the AlphaFold 3 server (50). Structures were visualized and aligned using PyMOL (version 2.5.7) and ChimeraX (version 1.10). Multiple sequence alignments were performed with Clustal Omega (89). Graphs were generated using Prism, and final figures were assembled in Adobe Illustrator (version 28.0.0.88).

## Data, materials, and software availability

All study data are included in the article and its supplementary materials. DNA constructs and transgenic parasite lines used in this study are available upon reasonable request.

## ACKNOWLEDGEMENTS

We thank Mathieu Brochet and Dominique Soldati-Favre from the Université de Genève for access to the research infrastructures required for cellular bioenergetics analysis, and Paul-Christian Burda from the Justus Liebig University Giessen for providing plasmids used in this study. Ac-H-FluNox was kindly provided by Tasuku Hirayama from Gifu Pharmaceutical University. We also thank Monja Paasche from the Bernhard Nocht Institute for Tropical Medicine for technical assistance, and Daisey Chen from University of California, San Diego, for whole-genome sequencing. This work was funded by the Bundesministerium für Forschung, Technologie und Raumfahrt (BMFTR; Federal Ministry of Research, Technology and Space; grant number 01KI2104 to JMM). Responsibility for the content of this publication lies solely with the authors. We also acknowledge financial support from the priority program SPP 2225 “Exit Strategies of Intracellular Pathogens” of the Deutsche Forschungsgemeinschaft (DFG; German Research Foundation; projects 531646514 to JMM and 446605368 to UD). LS is supported by a stipend from the Jürgen Manchot Stiftung. CB is the recipient of a Swiss Postdoctoral Fellowship from the Schweizer Nationalfonds (Swiss National Science Foundation, project 217028). JK receives funding from a generous donor advised by CARIGEST SA, acquired by Dominique Soldati-Favre. MRR was supported by an F32 postdoctoral fellowship from the National Institutes of Health (NIH; project F32AI188627).

## AUTHOR CONTRIBUTIONS

Conceptualization: DEG, JMM

Methodology: LS, MRR, JK, TZ, KSH, UD, JMM

Investigation: LS, MRR, JK, TZ, YS, ALS, RDE, KSH, CB

Data Curation and Analysis: LS, MRR, JK, TZ, UD, JMM

Funding Acquisition: LS, UD, CB, JMM

Writing – Original Draft: LS, MRR, JMM

Writing – Review & Editing: all authors

## SUPPLEMENTARY METHODS

### Filter-aided sample preparation

For proteome analysis, samples were prepared using a modified filter-aided sample preparation, as previously described (90, 91). Briefly, pelleted parasites were lysed in 200 µL of buffer containing 7 M urea, 2 M thiourea, 5 mM DTT, and 2% 3-[(3-cholamidopropyl)dimethylammonio]-1-propanesulfonate, followed by gentle vortexing and sonication at 4°C for 15 minutes. Lysates were centrifuged at 16,200 x g for 15 minutes at 4°C, and the supernatants were transferred into clean 1.5 mL tubes. Protein concentration was determined using the Pierce 660 nm protein assay (Thermo Fisher Scientific). 20 µg of protein were applied to Micron-30 spin filter columns (Merck) and washed three times with 8 M urea. Following reduction with DTT and alkylation with iodoacetamide (IAA), excess IAA was quenched with DTT, and the membrane was washed three times with 50 mM NH4HCO3. Proteins were digested overnight at 37°C with trypsin (Promega) at a 1:50 enzyme-to-protein ratio. Peptides were recovered by centrifugation and two additional washes with 50 mM NH4HCO3, acidified with 1% trifluoroacetic acid, lyophilized, and reconstituted in 0.1% formic acid for LC-MS analysis.

### Liquid chromatography – mass spectrometry

Proteome samples were analyzed using a nanoElute liquid chromatography system coupled online to a timsTOF Pro mass spectrometer (both from Bruker). Peptides corresponding to 200 ng were loaded in direct injection mode at 600 bar onto a reversed-phase C18 column (Aurora ULTIMATE UHPLC emitter, 25 cm x 75 µm, 1.7 µm particle size, IonOpticks) and separated at 400 nL/minute at 50°C. Mobile phase A was 0.1% formic acid in water, and mobile phase B was 0.1% formic acid in acetonitrile. Separation was achieved using a linear gradient from 2% to 37% B over 39 minutes, followed by a 5-minute rinse at 95% B.

Eluting peptides were analyzed in positive-mode ESI-MS using parallel accumulation serial fragmentation enhanced data-independent acquisition (diaPASEF) (92). The dual TIMS was operated at a fixed duty cycle close to 100%, with equal accumulation and ramp times of 100 ms, each spanning a mobility range from 1/K0 = 0.6 to 1.6 Vs cm^-2^. We defined 36 x 25 Th isolation windows from *m/z* 300 to 1,165, resulting in 2-3 diaPASEF scans per TIMS cycle and an overall cycle time of 1.7 seconds. Collision energy was ramped linearly as a function of mobility from 59 eV at 1/K0 = 1.3 Vs cm^-2^ to 20 eV at 1/K0 = 0.85 Vs cm^-2^. Samples were measured in triplicate.

### Raw data processing

Raw mass spectrometry data were processed using DIA-NN (version 2.2.0) (93) with default settings for library-free database search. A custom database was compiled containing the UniProtKB entries of the *P. falciparum* reference proteome and common contaminants (March 2025 release, 5,495 entries). Trypsin was specified as the protease for peptide identification and *in silico* library generation, allowing one missed cleavage. Carbamidomethylation was set as a fixed modification with no variable modifications allowed. Peptides of 7-30 amino acids were considered, with a precursor *m/z* range of 300-1,800 and a product ion *m/z* range of 200-1,800. For quantification, we used “QuantUMS (high precision)” mode with RT-dependent median-based cross-run normalization. Mass accuracy and scan window parameters for MS1 and MS2 were automatically optimized by DIA-NN. Peptide precursor false discovery rates were controlled below 1%.

---

**Supplementary Figure S1:**
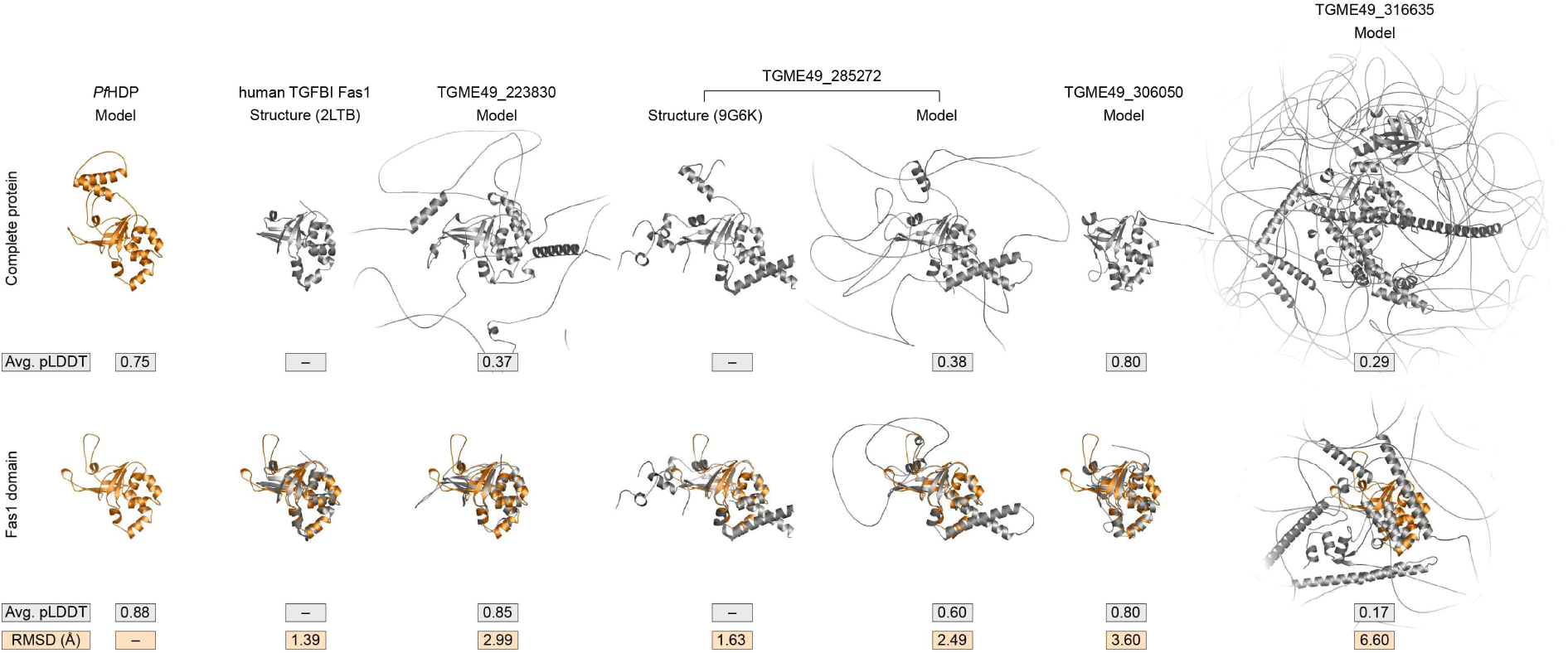
Structural alignment of HDP with Fas1-domain-containing proteins from *Toxoplasma gondii*. Shown are AlphaFold3 models of HDP and all Fas1-domain-containing proteins encoded in the *T. gondii* genome, along with the experimentally determined structures of the mitoribosomal subunit TGME49_285272 (22) and the human TGFBI Fas1 domain (51). Whole-protein structures are displayed in the top row, while the bottom row shows alignments of Fas1 domains only. Average pLDDT values for individual models and RMSD values for the alignments are indicated.

**Supplementary Figure S2:**
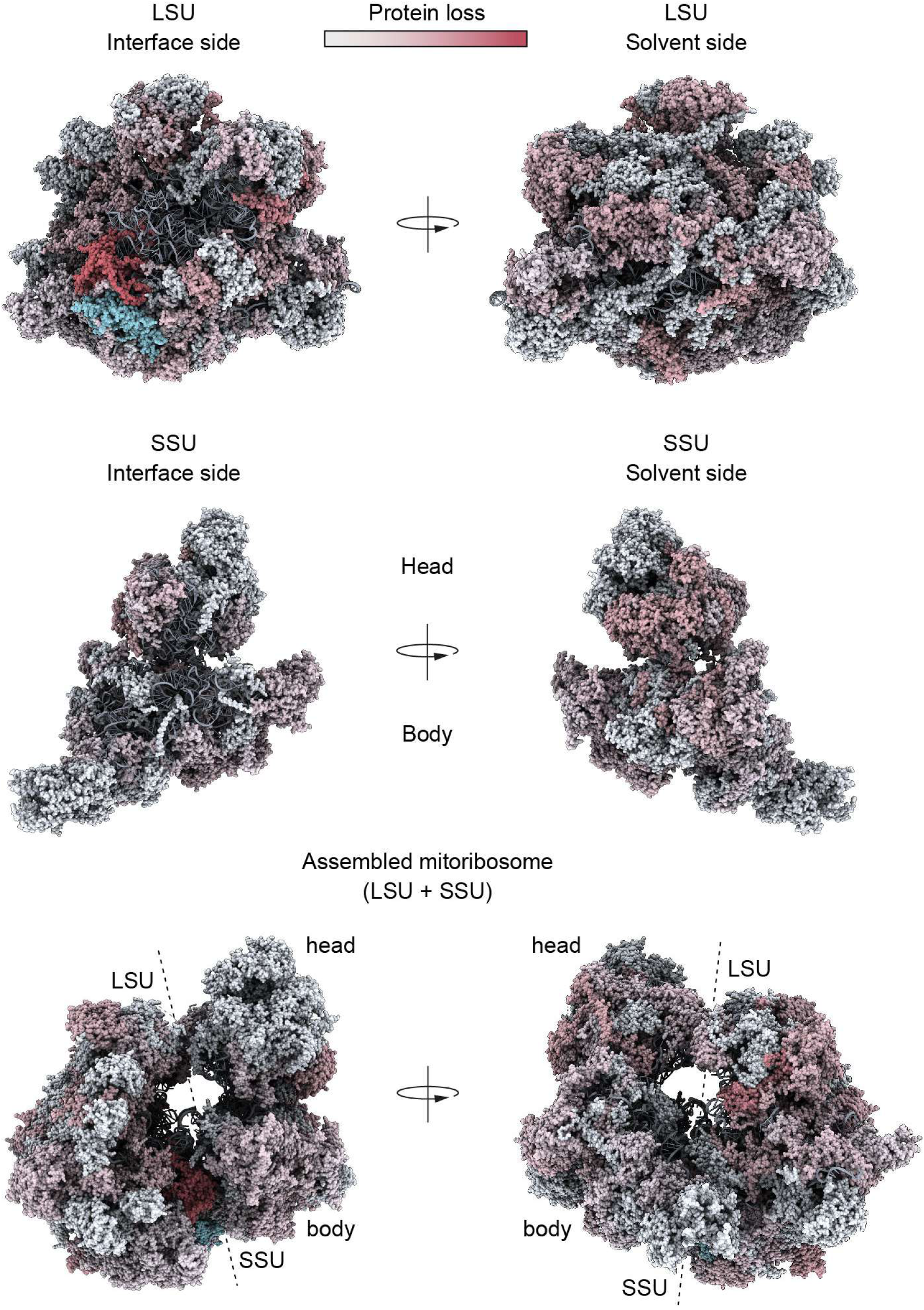
HDP-dependent reduction of *P. falciparum* mitoribosomal proteins mapped onto the *T. gondii* mitoribosome structure. Proteins are color-coded to indicate the reduction of their respective *P. falciparum* orthologs following HDP knockout. The putative HDP ortholog TGME49_285272 is highlighted in cyan. The large (LSU) and small (SSU) ribosomal subunits are shown from both the interface and solvent sides, with additional side-views of the fully assembled mitoribosome.

**Supplementary Figure S3:**
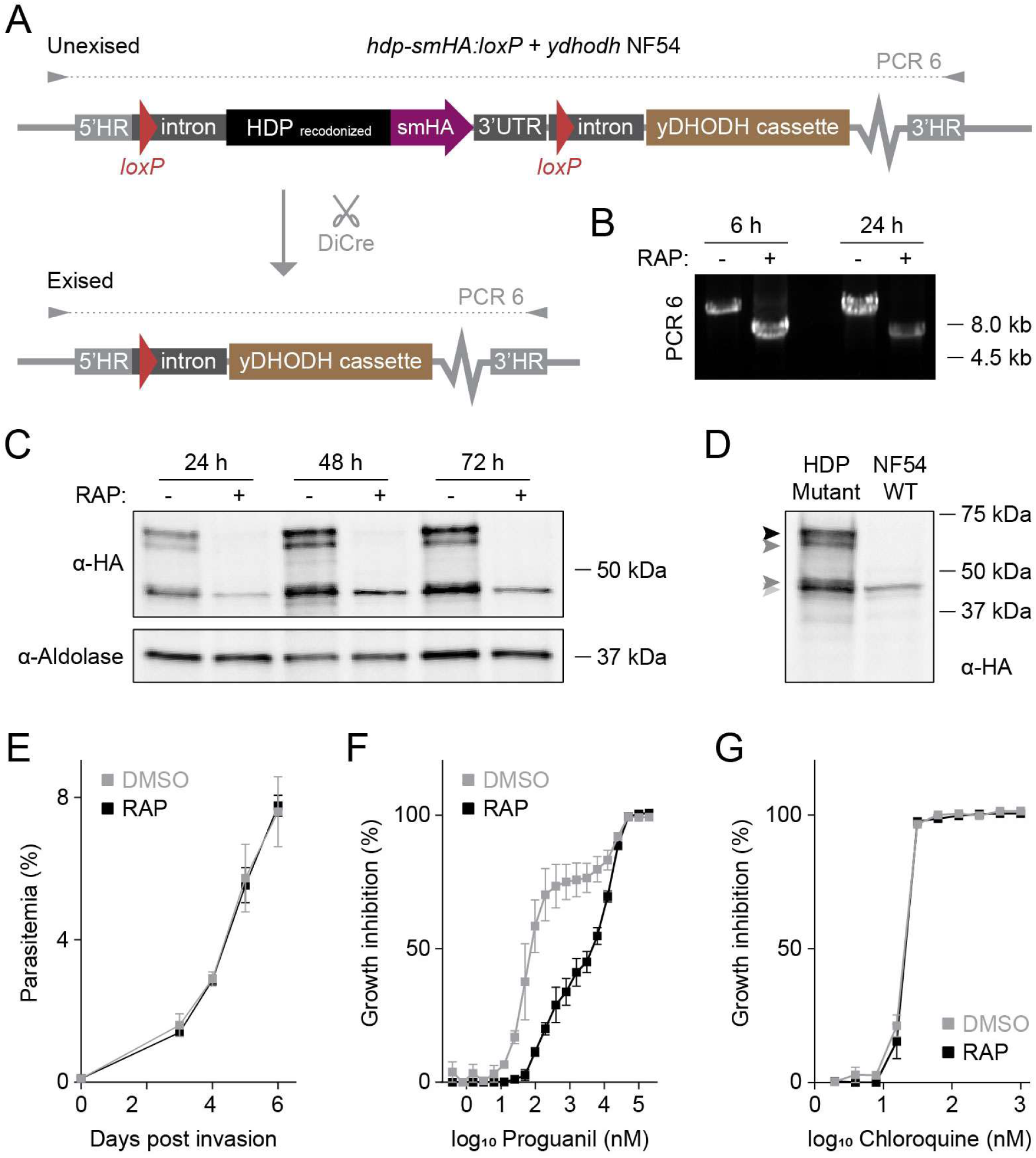
Generation and validation of an NF54-based conditional HDP knockout line metabolically rescued by yDHODH expression. **(A)** Genetic strategy for tagging and conditional deletion of full-length HDP in *P. falciparum* NF54 parasites. Shown is the recombined locus before and after DiCre-mediated excision. Primer combinations used for diagnostic PCR 6 are indicated. A yDHODH expression cassette is included to mediate metabolic rescue. **(B-D)** RAP-induced loss of HDP, confirmed by (B) diagnostic PCR 6 and (C) Western blot analysis. Time points for DNA and protein extraction are indicated. A non-specific protein band remains after RAP treatment and is also observed in NF54 control parasites (D). Black arrowhead: predicted size of HDP-smHA; dark grey arrowheads: possible processing products of either HDP or the smHA tag; light grey arrowhead: non-specific band. **(E)** *hdp-smHA:loxP + ydhodh* NF54 parasites exhibit normal growth in the presence of RAP. Shown are mean parasitemia +/- SEM. N = 3 independent experiments. **(F, G)** RAP treatment hypersensitizes *hdp-smHA:loxP + ydhodh* NF54 parasites to proguanil (F), but not to chloroquine (G). Growth-response curves are shown with mean +/- SEM values. N ≥ 3 independent experiments.

**Supplementary Figure S4:**
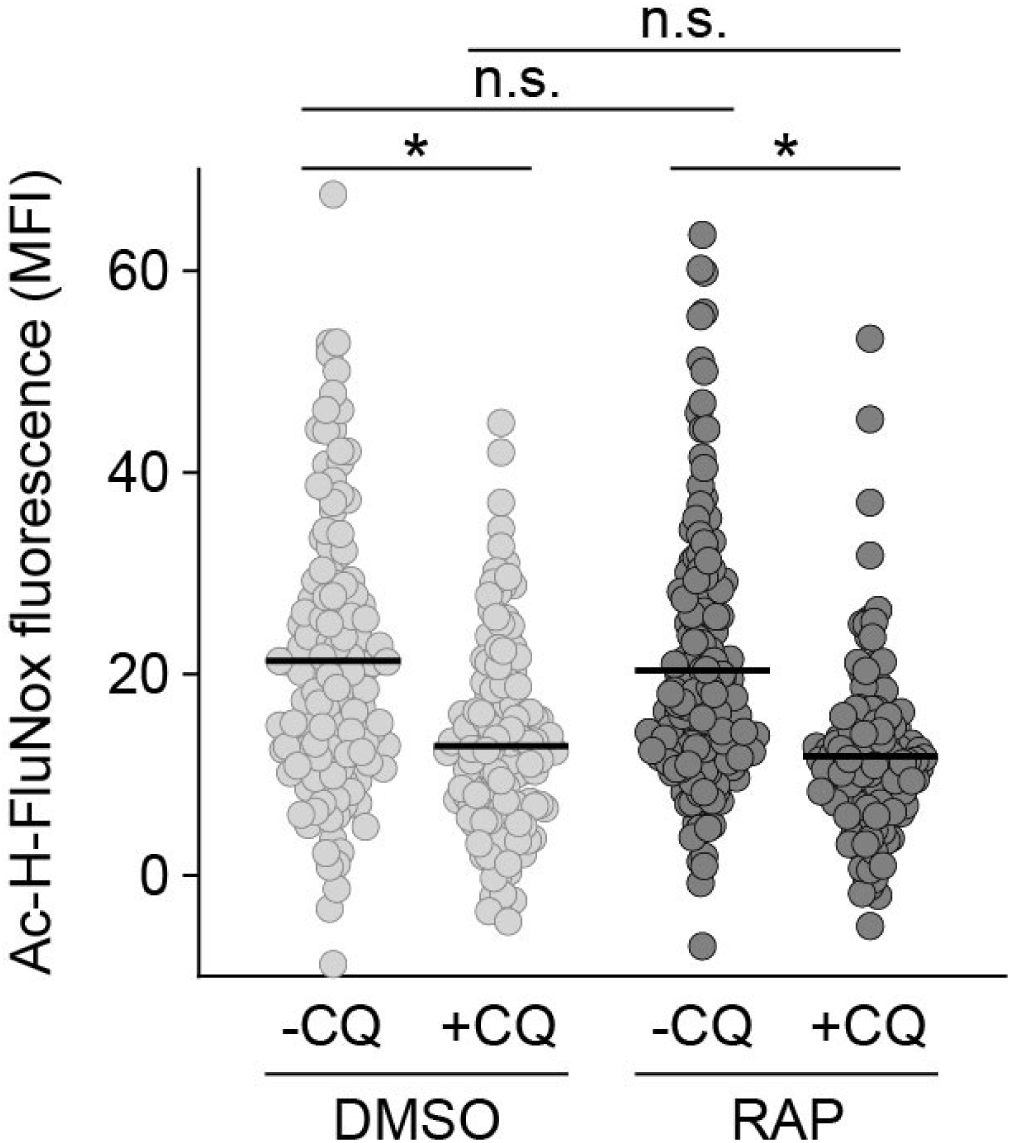
Loss of HDP does not affect levels of free cytosolic heme. DMSO and RAP-treated *hdp-smHA:loxP + ydhodh* NF54 parasites were incubated with or without chloroquine (CQ), stained with the labile heme probe Ac-H-FluNox, and analyzed by live-cell microscopy to determine mean fluorescence intensity (MFI). Shown are individual and mean values. n.s., non-significant; *P<0.05, one-way ANOVA with Tukey’s post hoc test; N ≥ 136 parasite from 5 independent experiments.

**Supplementary Table S1:**
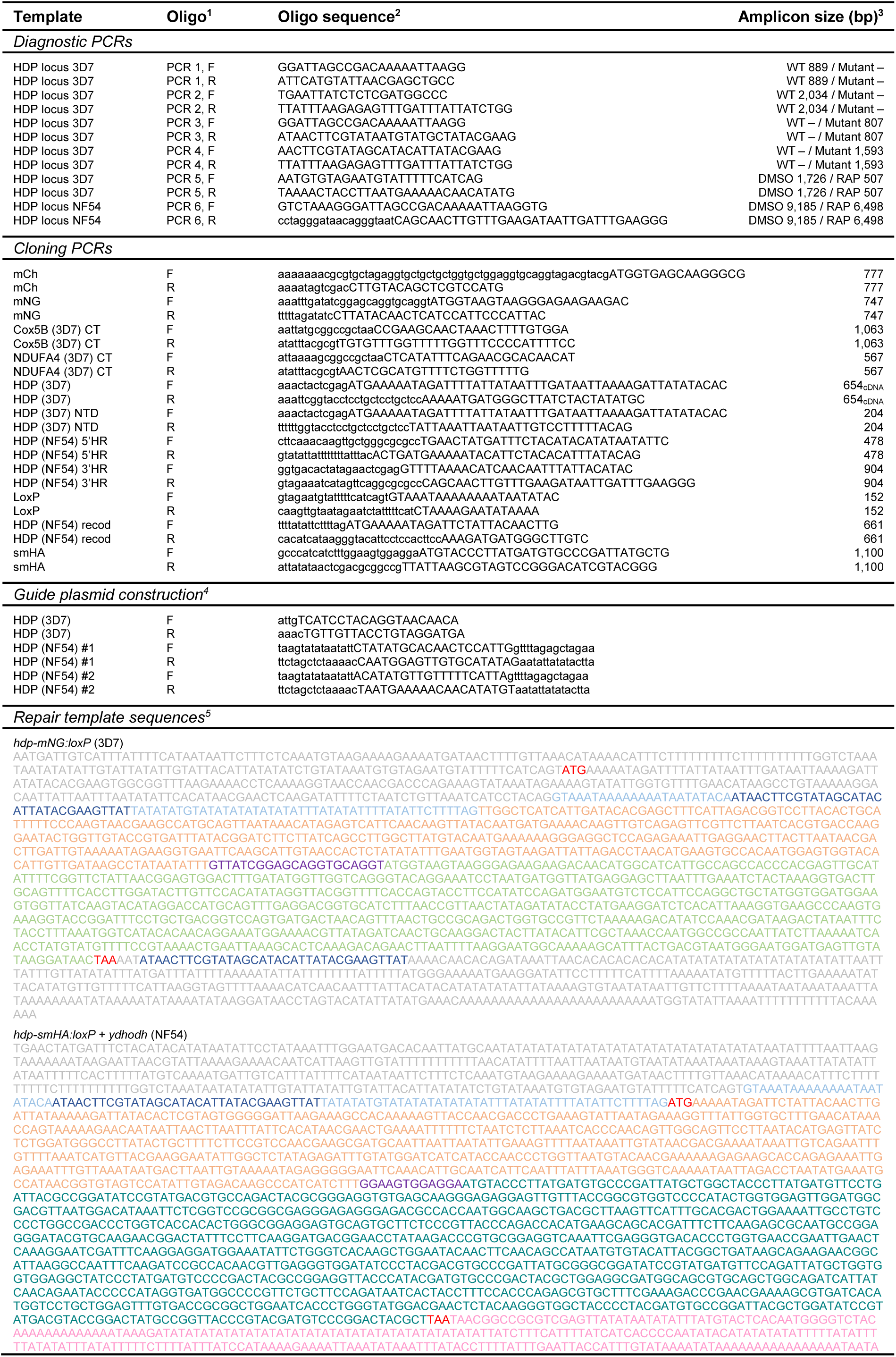
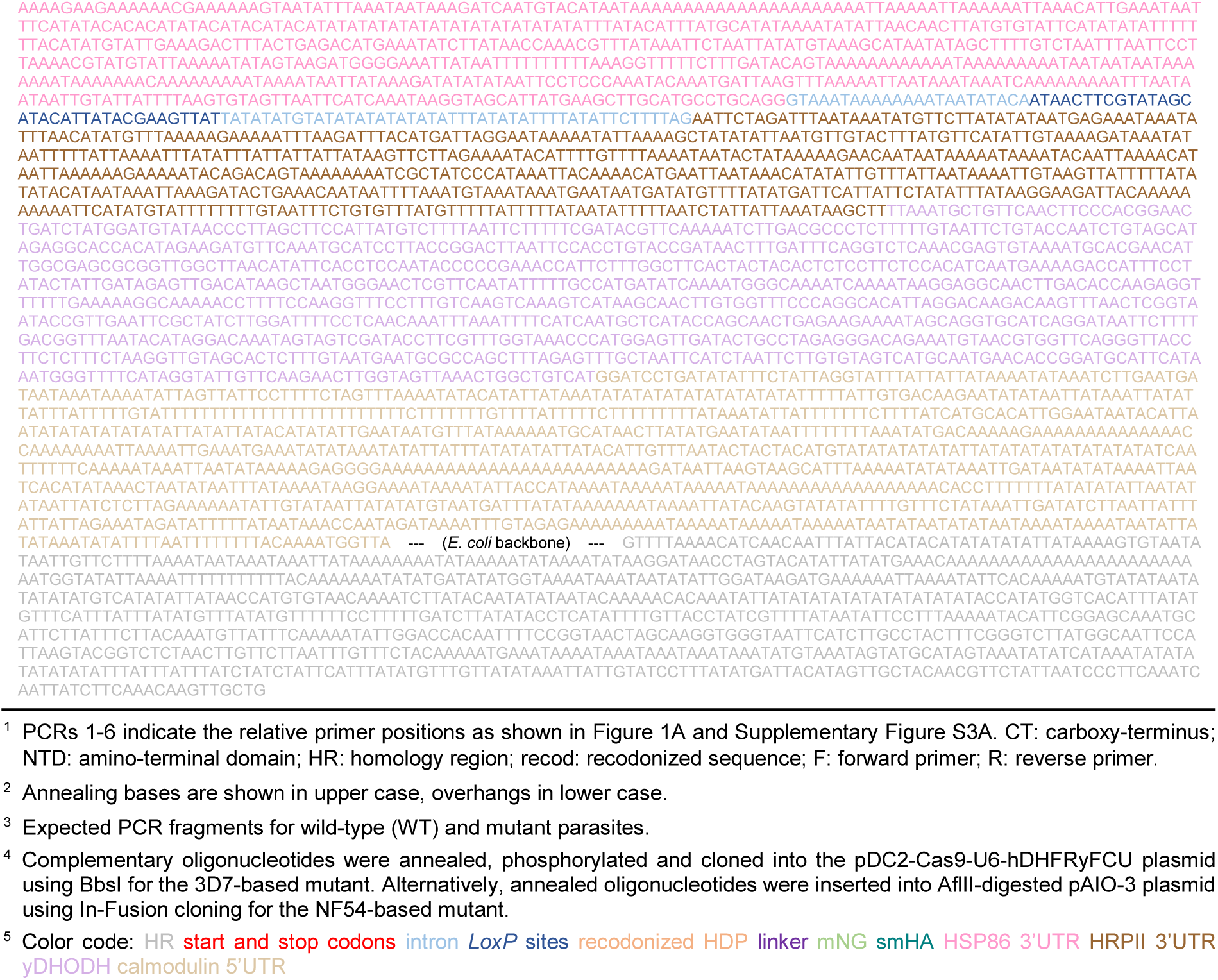
Oligonucleotides and repair templates used in this study.

**Supplementary Table S2:**
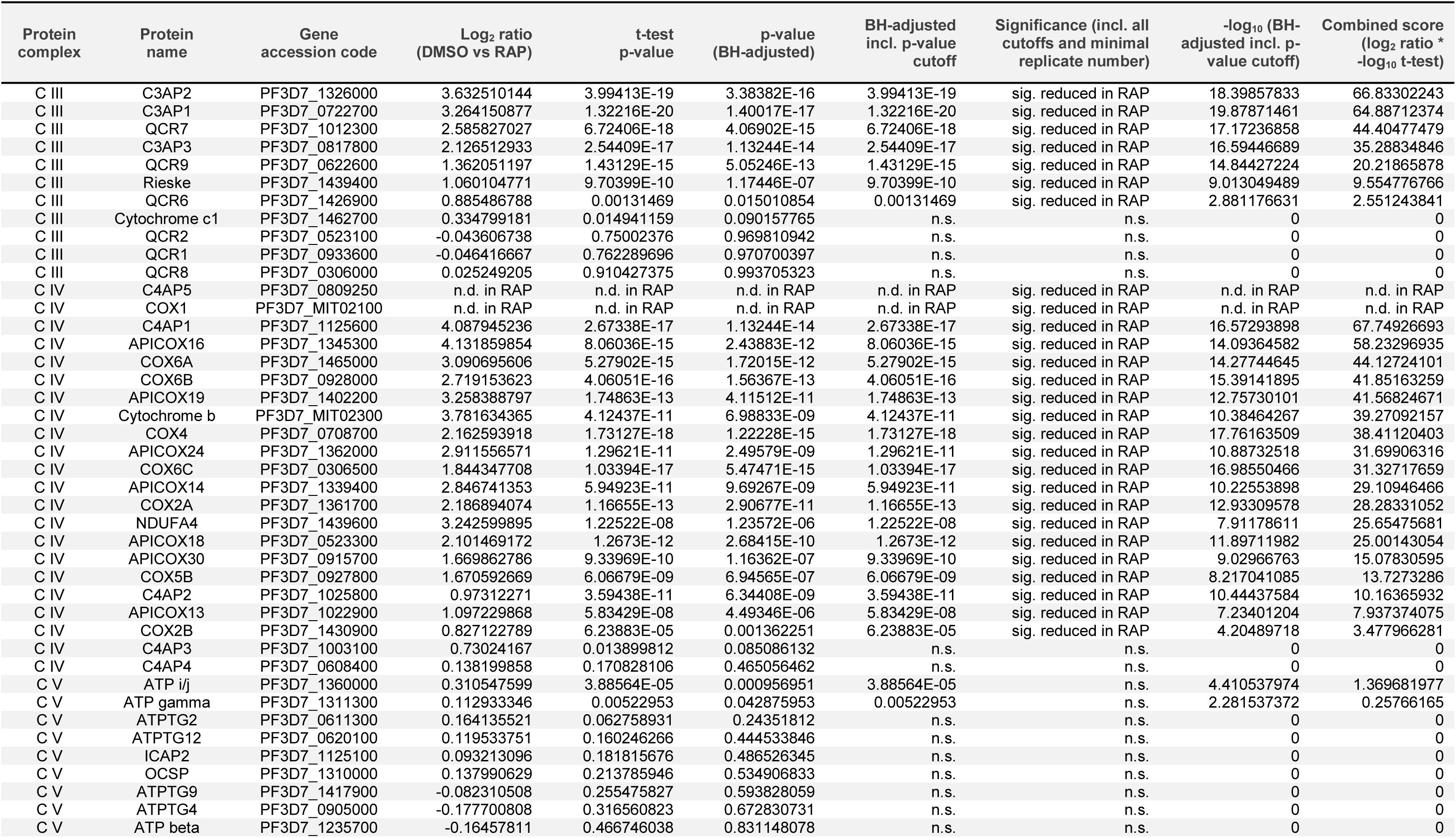
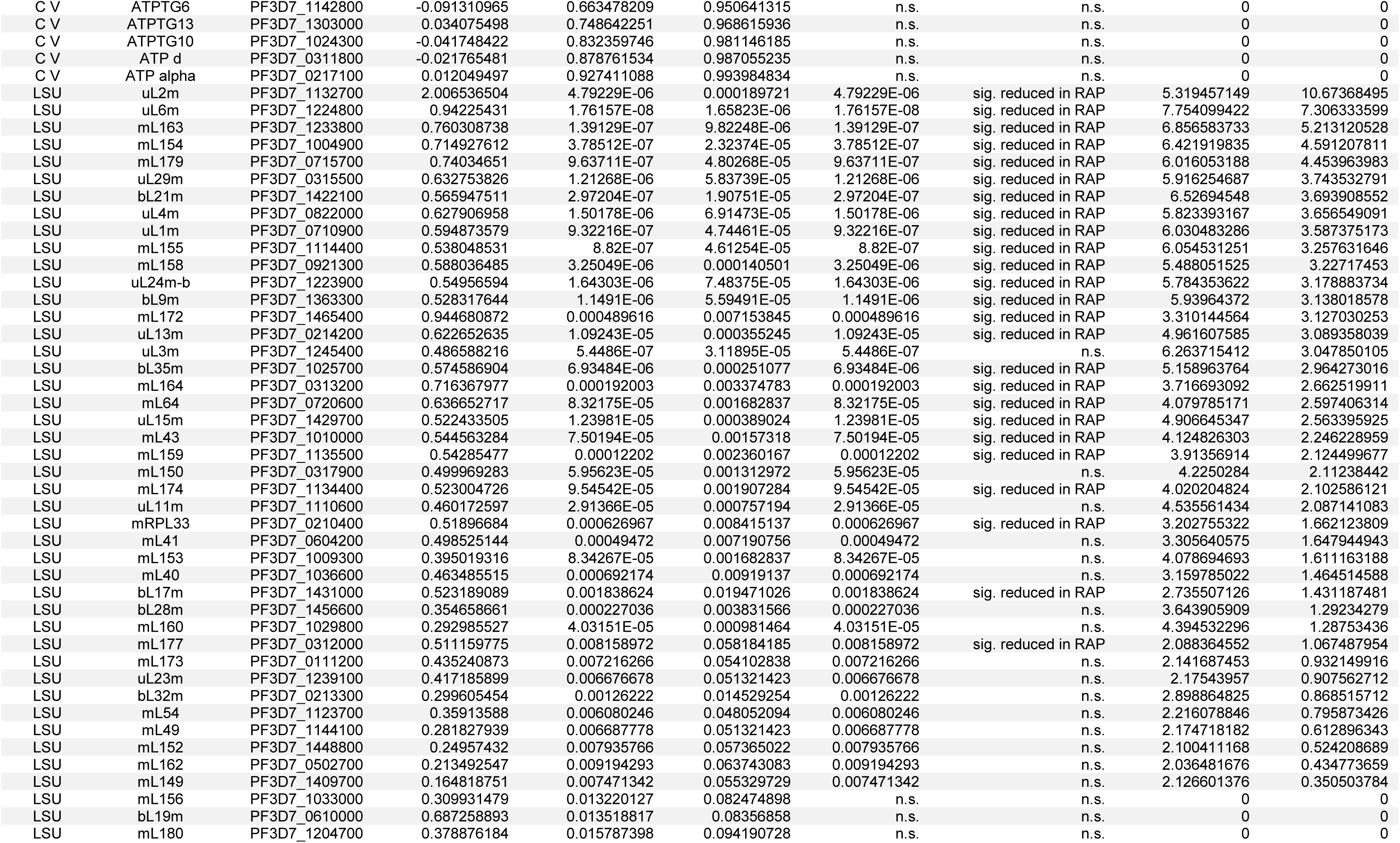
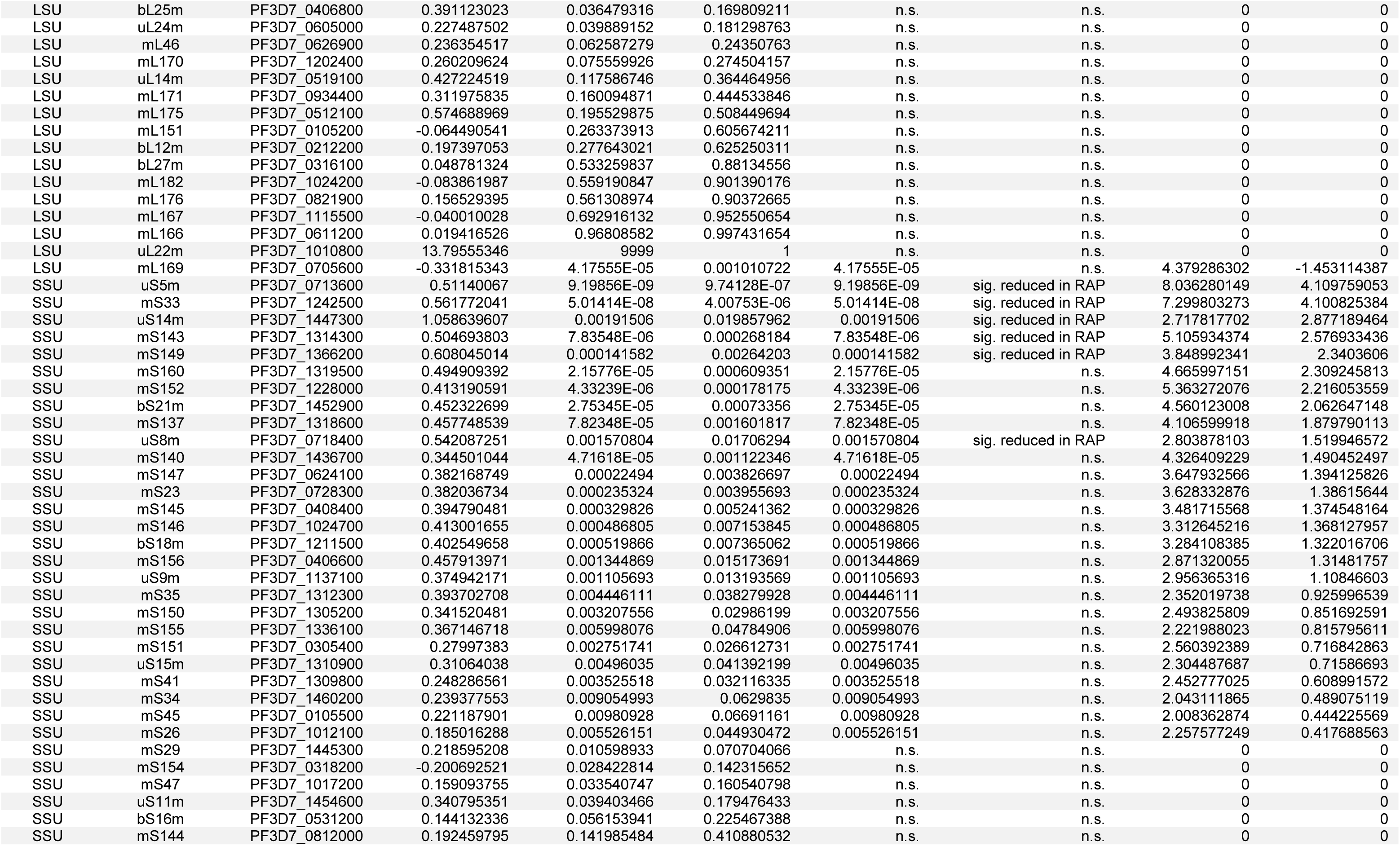
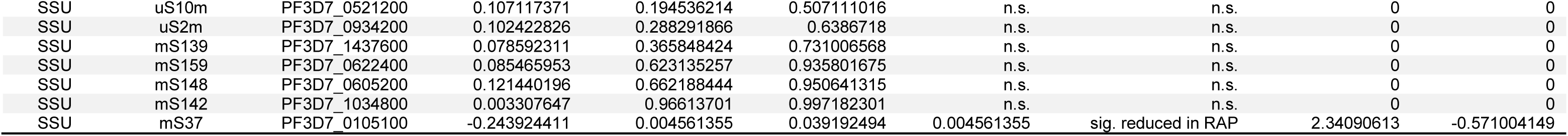
Quantitative proteomic analysis of respiratory chain complexes and mitoribosomal subunits in *hdp-mNG:loxP + ydhodh* parasites.

**Supplementary Table S3:**
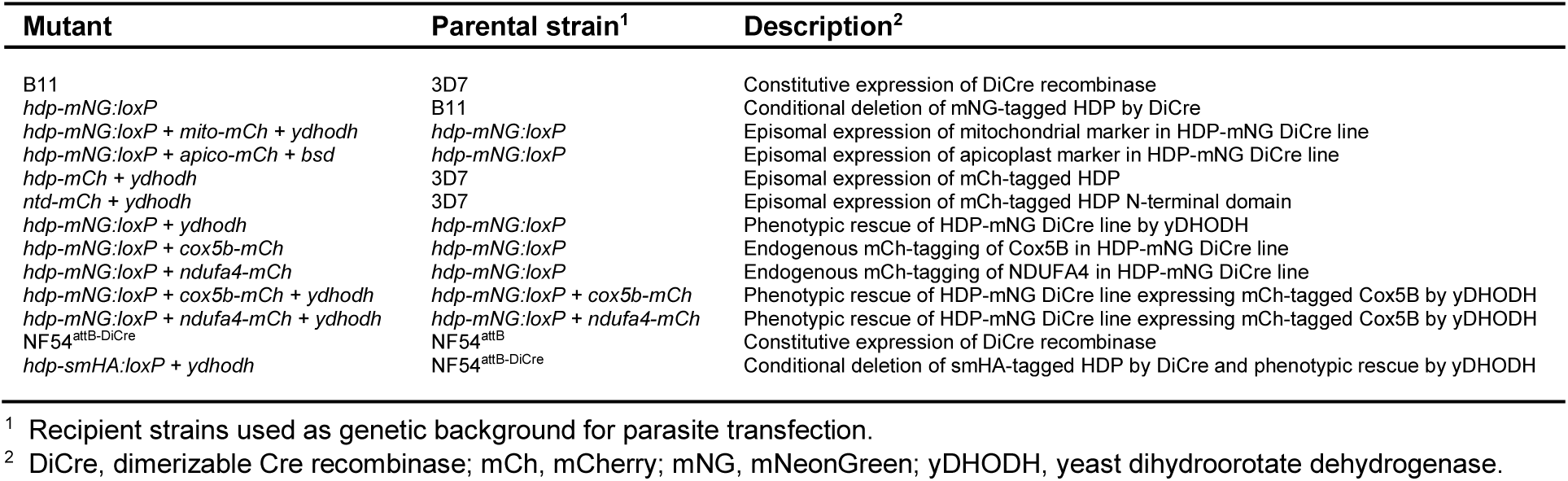
Transgenic parasite strains used in this study.

